# Insecticide resistance management strategies for public health control of mosquitoes exhibiting polygenic resistance: a comparison of sequences, rotations, and mixtures

**DOI:** 10.1101/2022.10.28.514211

**Authors:** Neil Philip Hobbs, David Weetman, Ian Michael Hastings

## Abstract

Malaria control uses insecticides to kill *Anopheles* mosquitoes. Recent successes in malaria control are threatened by increasing levels of insecticide resistance (IR), requiring insecticide resistance management (IRM) strategies to mitigate this problem. Field trials of IRM strategies are usually prohibitively expensive with long timeframes, and mathematical modelling is often used to evaluate alternative options. Previous IRM models in the context of malaria control assumed IR to have a simple (monogenic) basis, whereas in natural populations, IR will often be a complex polygenic trait determined by multiple genetic variants.

A quantitative genetics model was developed to model IR as a polygenic trait. The model allows insecticides to be deployed as sequences (continuous deployment until a defined withdrawal threshold, termed “insecticide lifespan”, as indicated by resistance diagnosis in bioassays), rotations (periodic switching of insecticides), or full-dose mixtures (two insecticides in one formulation). These IRM strategies were compared based on their “strategy lifespan” (capped at 500 generations). Partial rank correlation and generalised linear modelling was used to identify and quantify parameters driving the evolution of resistance. Random forest models were used to identify parameters offering predictive value for decision-making.

Deploying single insecticides as sequences or rotations usually made little overall difference to their “strategy lifespan”, though rotations displayed lower mean and peak resistances. Deploying two insecticides in a full-dose mixture formulation was found to extend the “strategy lifespan” when compared to deploying each in sequence or rotation. This pattern was observed regardless of the level of cross resistance between the insecticides or the starting level of resistance. Statistical analysis highlighted the importance of insecticide coverage, cross resistance, heritability, and fitness costs for selecting an appropriate IRM strategy.

Full-dose mixtures appear the most promising of the strategies evaluated, with the longest “strategy lifespans”. These conclusions broadly corroborate previous results from monogenic models.

**Author Summary:** Insecticides impregnated into bed-nets or sprayed on walls are used to kill the *Anopheles* mosquitoes which transmit malaria. Unfortunately, the usage of insecticides has inevitably led to mosquitoes evolving resistance to the toxic effect of these insecticides. Insecticide resistance management strategies may be used to slow the rate of resistance evolution, however which strategies are effective, and when they are effective, is often unclear. Previous models evaluating insecticide resistance management strategies have assumed resistance is encoded by a single gene (is a monogenic trait). However, in natural populations resistance may be determined by multiple genes (is a polygenic trait). It is unclear whether such increased model complexity may change predictions We modelled resistance as a polygenic trait and found little difference in the benefit between rotating insecticides regularly versus deploying continuously until resistance reaches a critical threshold then switching. In contrast, mixtures combining two insecticides extended the projected lifespan of the insecticides, even when they share resistance mechanisms (cross resistance). Similar findings from previous monogenic models, strengthen support for the use of full-dose mixtures.

## Introduction

Long-lasting insecticide-treated nets (LLINs) and indoor residual spraying (IRS) play a dominant role in reducing the burden of malaria (Bhatt et al., 2015). The evolution of insecticide resistance (IR) poses a major threat to sustained reductions in malaria transmission and prevalence (e,g, Hemingway et al., 2016). Pyrethroids are the primary insecticide class used on LLINs, but pyrethroid resistance is geographically widespread (Hancock et al., 2020) and is increasing in frequency and intensity (Ranson & Lissenden, 2016). How entomological outcomes (i.e., greater mosquito survival) translate into impacts on malaria epidemiology is uncertain and complex (Van Hul et al., 2021). However, tools designed to kill mosquitoes or prevent them from blood-feeding, which cease to be capable of doing so as a result of resistance (Asidi et al., 2012; Irish et al., 2008), are clearly suboptimal.

The Global Plan for Insecticide Resistance Management (GPIRM) was developed to help mitigate the potential impact IR (WHO, 2012). Compared to agriculture, public health has relatively few insecticide classes available (Ranson & Lissenden, 2016). Insecticides for malaria control are mainly deployed to target adult mosquitoes indoors as either LLINs or IRS, requiring the insecticides to be safe for close human contact, which constrains the development of novel products. LLINs are designed to have 3- year lifespans, whilst IRS is usually deployed annually (WHO, 2020). Therefore, insecticides used in an insecticide resistance management (IRM) strategy for public health cannot be changed rapidly, unlike agriculture where re-application can occur monthly or even weekly. This necessity for safe and long-lasting insecticides has led to an over-reliance on a small number of insecticides, predominantly pyrethroids (Oxborough, 2016).

There is a need to evaluate IRM strategies designed to prolong the operational lifespan of insecticides to ensure future insecticides are not rapidly lost to resistance. This is especially important considering the new insecticides being developed by the Innovative Vector Control Consortium and industry partners. Evaluating IRM strategies in the laboratory and field is challenging due to the number of potential IRM strategies, the need for replication over different ecological/epidemiological settings, and the long study durations needed to observe phenotypic changes in resistance. Cluster randomised control trials used for evaluating the epidemiological effectiveness of malaria vector control tools generally last no longer than 2-3 years (Wilson et al., 2015), which is unlikely to be of sufficient duration to detect the benefit of any one IRM strategy over another. Mathematical modelling and computer simulations can simulate the IR response to IRM strategies over decades, and therefore provide a valuable evaluation tool (Tabashnik, 1986). Computer simulations can be run over numerous scenarios to provide insights into which IRM strategies are optimal/sub-optimal under different ecological, epidemiological, and operational contexts. Evaluating IRM strategies using computer simulations allows for the identification of potential problems with IRM strategies prior to implementation in the field where expensive and time-consuming errors can be made.

Theoretical and mathematical modelling of IRM for agriculture and public health has a long history (e.g. Comins, 1977; Curtis, 1985; Mani, 1985; Wood, 1981). However much of this previous work has modelled IR as a monogenic trait i.e., a trait encoded by a single mutation in a single gene. Few modelling studies have considered IR to be a polygenic trait (but see, for example, Gardner et al., 1998; Haridas & Tenhumberg, 2018; Via, 1986), of which none have evaluated IRM strategies in a public health context. In more recent years, mathematical models have been developed which take advantage of increased computational power allowing for improved evaluation of IRM strategies in agriculture (e.g., Helps et al., 2017) and public health (e.g., Levick et al., 2017; Madgwick & Kanitz, 2022, Hastings et al., 2022), of which these examples are monogenic models.

Public health IRM recommendations from models are therefore currently limited to scenarios where resistance in the field is monogenic. However mosquitoes are known to harbour a diverse array of resistance genes and mechanisms, including target site, metabolic, cuticular and behavioural resistance mechanisms (Balabanidou et al., 2018), alongside non-specific, largely environmentally influenced factors associated with increased tolerance to insecticides, for example mosquito age, and physiological condition (Lissenden et al., 2021)

Resistance management strategies often have confusing naming systems, with the same resistance management strategies called multiple names. This potentially confusing terminology has resulted from the often-independent development and evaluation of resistance management strategies across disciplines (Peck, 2001; Rex Consortium, 2007). Therefore, we define our terminology here for the three IRM strategies that will be investigated, and how this relates to terms used in previous work:

- Sequences, involve continuously deploying a particular insecticide formulation until a designated level of resistance is reached (the withdrawal threshold), at which point it is replaced by another insecticide to which there is less resistance. This strategy is also referred to as responsive alternation or series application in the literature (Rex Consortium, 2013). The threshold for withdrawing the current insecticide and replacing with the next insecticide the sequence is usually based on a bioassay survival threshold or, in the case of monogenic traits, an allele frequency threshold.
- Rotations, also referred to as cycling, alternation, or periodic application (Rex Consortium, 2013), involve the periodic pre-planned switching between insecticide formulations over time, such that one insecticide is temporarily replaced with another insecticide.
- Mixtures are confusingly also occasionally referred to as combinations (Rex Consortium, 2013) or pyramiding when referring to transgenic crops (Roush, 1998). In public health, the term combination would more generally refer to the use of two different insecticides within the same household, but not in the same control tool, such as deploying both LLINs and IRS (WHO, 2012). We define mixtures as the use of two insecticides in a single formulation such that the target insect inevitably encounters both insecticides simultaneously.

A comprehensive review of IRM modelling highlighted a lack of models including quantitative resistance and cross resistance (Rex Consortium, 2010). Quantitative genetic modelling assumes many genes determine IR level and each gene has only a minor phenotypic effect. It is the accumulated effect of these genes which combine to give larger phenotypic effects (Walsh & Lynch, 2018). In this study we model IR as a quantitative trait, assuming a polygenic basis of resistance and allowing for cross-resistance between insecticides.

We present a flexible quantitative genetics model for the evaluation of IRM strategies. The model is calibrated primarily for *Anopheles gambiae* but can be adapted for other vector species. We investigate how insecticides could be deployed temporally, as either sequences, rotations, or mixtures. First, we compare the sequence and rotation strategies, i.e., where insecticides are deployed individually. Second, we compare sequences and rotations against mixtures, to understand the impact of deploying two insecticides in a single formulation. We apply a range of statistical methods to the data generated from the model to reveal insights into the drivers of the evolution of IR and highlight parameters which may be beneficial to measure in the field to improve the decision-making process for choosing the optimal IRM strategy for local conditions.

## Methods

### An Overview of the Quantitative Genetics Model

In the model described here, IR is assumed to be a classically quantitative genetic trait encoded by a large number of genes, each with very small effect on the phenotype (this assumption means genetic variances do not change substantially over selection). The model assumes discrete non-overlapping generations of mosquitoes, a standard assumption in both quantitative and population genetics modelling. Female mosquitoes lay eggs, which hatch, develop, and mature into adults. Adult mosquitoes of both sexes may then encounter the deployed insecticide(s). Female mosquitoes then lay the eggs that constitute the next discrete generation and then die, i.e., the model allows for only a single insecticide exposure per generation. We assume mating between male and female mosquitoes occurs after the insecticide encounters, i.e., after insecticide selection which is consistent with equivalent monogenic models (e.g. Levick et al., 2017; Madgwick & Kanitz, 2022), and that all females successfully mate. This ordering of insecticide exposure, mating, then dispersal is important as this impacts the level of selection (Sudo et al., 2018). The model tracks IR in two locations. First, the intervention site, where insecticidal interventions can be deployed, and operational decisions are made. Second, the refugia, where the insecticides under consideration are not deployed. We assume mating occurs within the respective locations i.e., intervention or refugia. Mated female mosquitoes can then migrate between the intervention site and “refugia”. Since mating occurs within the intervention site/refugia, all females successfully mate prior to migration and as females mate only once the model need not consider migration of males (because there would be no un- mated females available to mate with).

A list of our model assumptions can be found in Table 1.

**Table 1:** Assumptions for the Quantitative Genetics Model.

| **Table 1: Assumptions for the Quantitative Genetics Model** |  |  |
| --- | --- | --- |
| Here we outline our key model assumptions with an explanation of why they were made. |  |  |
| Number | Assumption | Rationale/Explanation |
| 1 | Mating in the population is random. | Standard assumption of quantitative genetics / population genetics modelling. |
| 2 | The population size is sufficiently large to prevent inbreeding depression. | Standard assumption of quantitative genetics modelling and likely to reflect reality for most wild mosquito populations. |
| 3 | Fitness costs are constant regardless of the level of insecticide resistance. | Fitness costs must occur within the Normal distribution. As the standard deviation stays constant the difference between the most and least resistant individuals remains constant. |
| 4 | Only a single generation is tracked at any one time point, which does not interbreed with older or younger generations. | Standard assumption of quantitative genetics modelling and population genetics modelling. |
| 5 | The relative population size is directly proportional to intervention | We define intervention coverage as the proportion of the mosquito |
|  | coverage in the intervention site and refugia. | population covered by the intervention. |
| 6 | Male insecticide exposure is proportional to, and generally less than, female insecticide exposure. | Female mosquitoes are the ones that blood feed and are therefore more likely to encounter insecticides used indoors. |
| 7 | Deployment decisions are made at the same generation as bioassays are conducted. | This is currently done for ease of computation. In the real-world bioassays would need to be done in sufficient time to allow for the results to be used to inform insecticide purchasing and deployment decisions. |
| 8 | Resistance mechanisms are limited to those which would affect bioassay survival. Behavioural resistance is not accounted for. | Behavioural resistance is likely to be based around mosquitoes reducing their exposure to the insecticide; and is therefore not readily measured in standard bioassays. |
| 8 | There is only dispersal between the intervention site and refugia. | Standard assumption in two-patch models (see Comins (1977)). |
| 10 | There are ten mosquito generations per year. | The number of generations can be updated if necessary for other insect species. This would then require a |
|  |  | recalculating of the Exposure Scaling Factor (Beta) to calibrate the simulations to the desired timescale. |
| 11 | There is always an insecticide in deployment. | We would expect constant coverage of interventions such as LLINs. The current model does not allow for gaps in deployment of insecticides. IRS deployments are more likely to be an on-off system; mainly being used to cover the main transmission seasons. |
| 12 | Decisions are made at the specified withdrawal and return thresholds. | Withdrawal of an insecticide does not happen until the threshold is reached (e.g., 10% survival). This means that if the survival at the next deployment interval is 9.9%, the insecticide will be redeployed. |
| 13 | Insecticides are deployed in the numerical sequence (1, 2, 3 etc). | The optimal insecticide (based on mosquito survival) is not necessarily the one deployed. |
| 14 | Insecticide survival in the field is directly correlated with bioassay survival. | Churcher et al (2016) found a linear relationship between bioassay mortality and mortality in experimental hut trials. |
| 15 | The deployment interval/frequency is fixed for a single simulation. The choice of what insecticide to deploy can only be made at that timepoint. | Interventions are likely to be distributed at fixed timepoints (e.g., IRS to cover the high transmission wet seasons). |
| 16 | The populations (intervention site and refugia) are isolated; there is no immigration/emigration outside these areas. | Standard quantitative/population genetic assumption. |
| 17 | Intervention site and refugia remain fixed sizes throughout the simulation | Model is deterministic. |
| 18 | Dispersal rates remain fixed throughout the simulation | Model is deterministic. |
| 19 | Heritability of resistance remains fixed throughout the simulation. | Model is deterministic. The heritability can be different for each insecticide. |
| 20 | Insecticides simulated are in public health used only and there are no non-public health insecticides deployed that may show cross resistance with the insecticides being simulated. | This means that all insecticide selection pressures are known and accounted for and makes the model more applicable for the evaluation of new insecticides. |
| 21 | Only a single mosquito species is present. | Operational decisions are made only on the resistance status of a single mosquito species which is often the case. |

### Developing a Quantitative Insecticide Resistance Scale

IR has many different definitions including survival in bioassays (WHO cylinder or CDC bottle), experimental huts, etc and there is no single clearly-defined scale that can be used to describe a “resistance phenotype” because this depends on the how the phenotype is measured. For example, survival in a bioassay may differ from survival in an experimental hut or insecticide sprayed wall in a house. To develop a polygenic model which allows for the tracking of quantitative IR, the first step is to develop an underlying scale of IR that can be converted to a measurable phenotype, in this case mosquito survival to insecticide exposure. For many quantitative traits this is a trivial process, where the trait of interest also corresponds to the amount of something produced, especially in agriculture and selective breeding where quantitative genetics is frequently leveraged. In contrast, resistance to insecticides is generally measured as a binary outcome as individuals either survive or die exposure.

In the field, the IR status of a mosquito population is typically measured using standardised diagnostic dose bioassays such as CDC bottle or WHO cylinder bioassays. These bioassays give the proportion of mosquitoes surviving contact with the insecticide (the measurable phenotype). We created a scale termed the Polygenic Resistance Score (PRS), denoted by *z*, which we use as a measure of resistance, and is described in more detail below. This underlying PRS scale produces a bioassay survival phenotype described by the Hill-variant of the Michaelis-Menten equation (where *n*=1) which converts the PRS to bioassay survival.

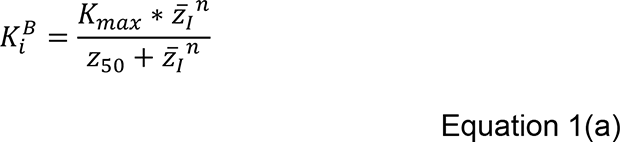

*K*^*B*^_*i*_ is the proportion of mosquitoes surviving in a diagnostic dose bioassay to insecticide *i* when they have a population mean PRS of *z̄* for corresponding resistance trait *I*. *K*_*max*_ is the maximum proportion of mosquitoes that could survive in a bioassay, which is 1. The *z*_50_ value is the PRS that gives 50% bioassay survival. For ease of interpretability, we have scaled the PRS such that *z̄*=100 equates to 10% bioassay survival. The *z*_50_ for this scale is therefore calculated from equation 1a to be 900. Our criterion for withdrawing an insecticide is >10% bioassay survival (<90% mortality) which is defined as the point when resistance is confirmed (WHO, 2018) so the evolution of resistance occurs over a scale of z=0 (the starting point) to z=100 (the withdrawal threshold). We note that our withdrawal threshold is set optimistically low, and in part this is done to help “force” one strategy to be better than other, if the withdrawal threshold is set high then all strategies would run to the end of the simulation and therefore all appear equally good. This relationship between PRS and bioassay survival is shown in Figure 1. This calibration of *K*_*max*_ = 1 and *z*_50_ = 900 is stable (i.e., bioassay mortality=10% when z=100) for population standard deviations up to 25 (Supplement 2, Table S2). A full description of all model parameters can be found in Table 2.

**Figure 1.**
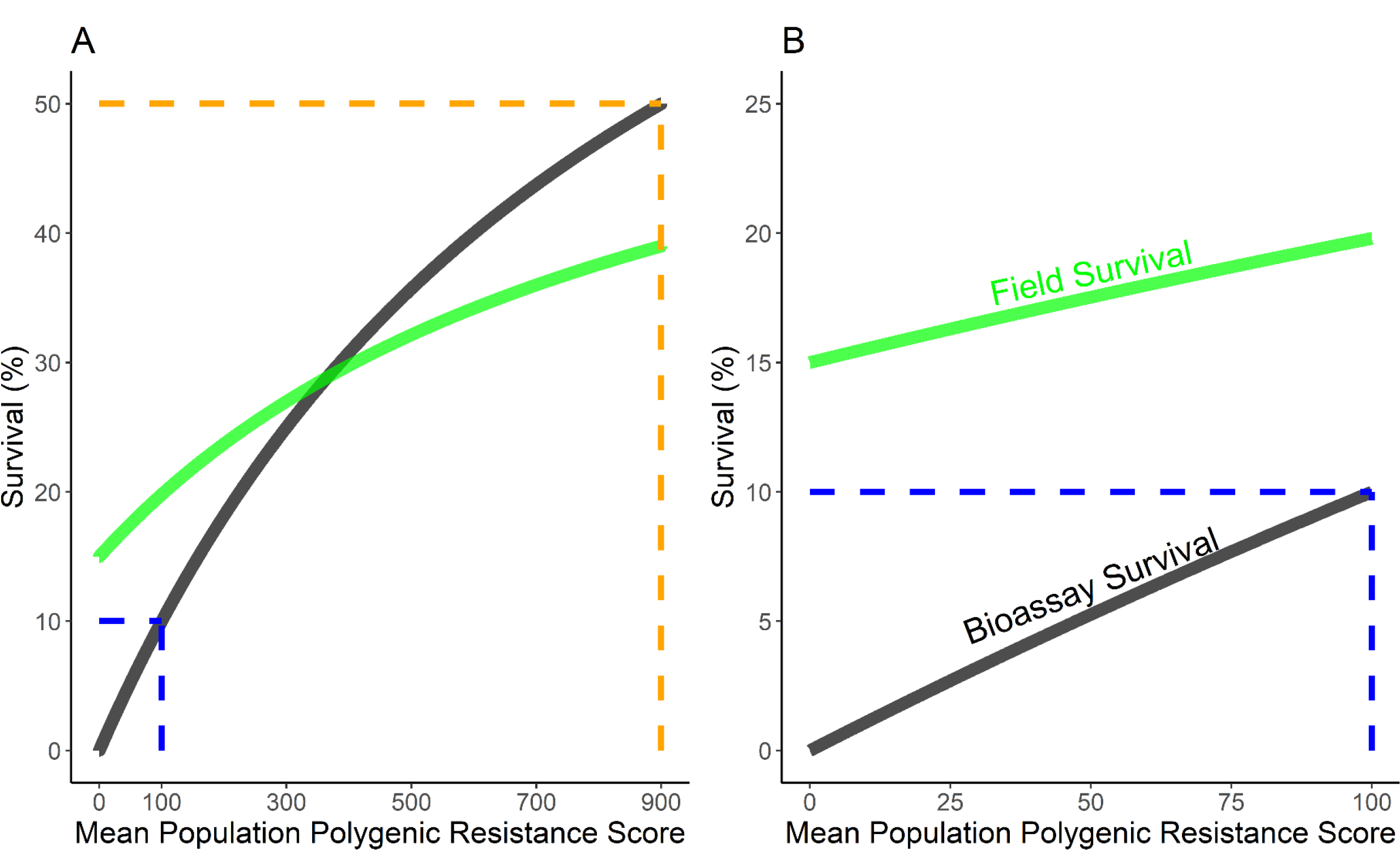
The relationship between the polygenic resistance score (PRS) and bioassay survival. The PRS was developed as a quantitative scale of insecticide resistance. The PRS is converted to bioassay survival as described in Equation 1a. The bioassay survival is converted to field survival based on a linear model of experimental hut survival predicted by WHO cylinder bioassay survival. The black line indicates the relationship between the PRS, and bioassay survival as calculated from Equation 1a. The green line indicates the relationship between the PRS, and field survival as calculated from Equation 1b. The dashed blue line is the withdrawal threshold (10% bioassay survival). The dashed orange line indicates the *z*_50_ of the polygenic resistance score scale, which was set at 900 (Panel A). Panel B is restricted to PRS values below our withdrawal threshold of PRS = 100.

**Table 2:**
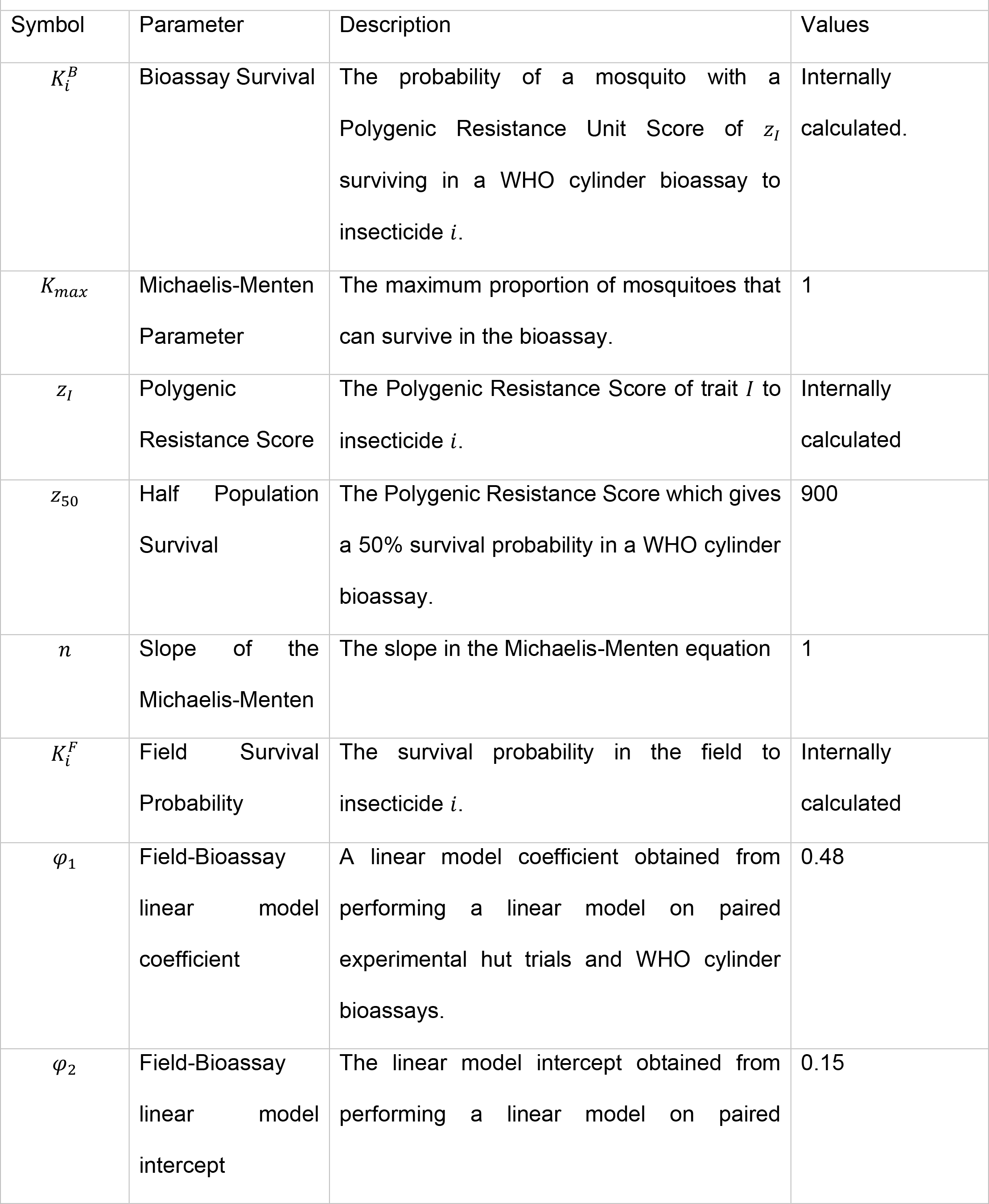
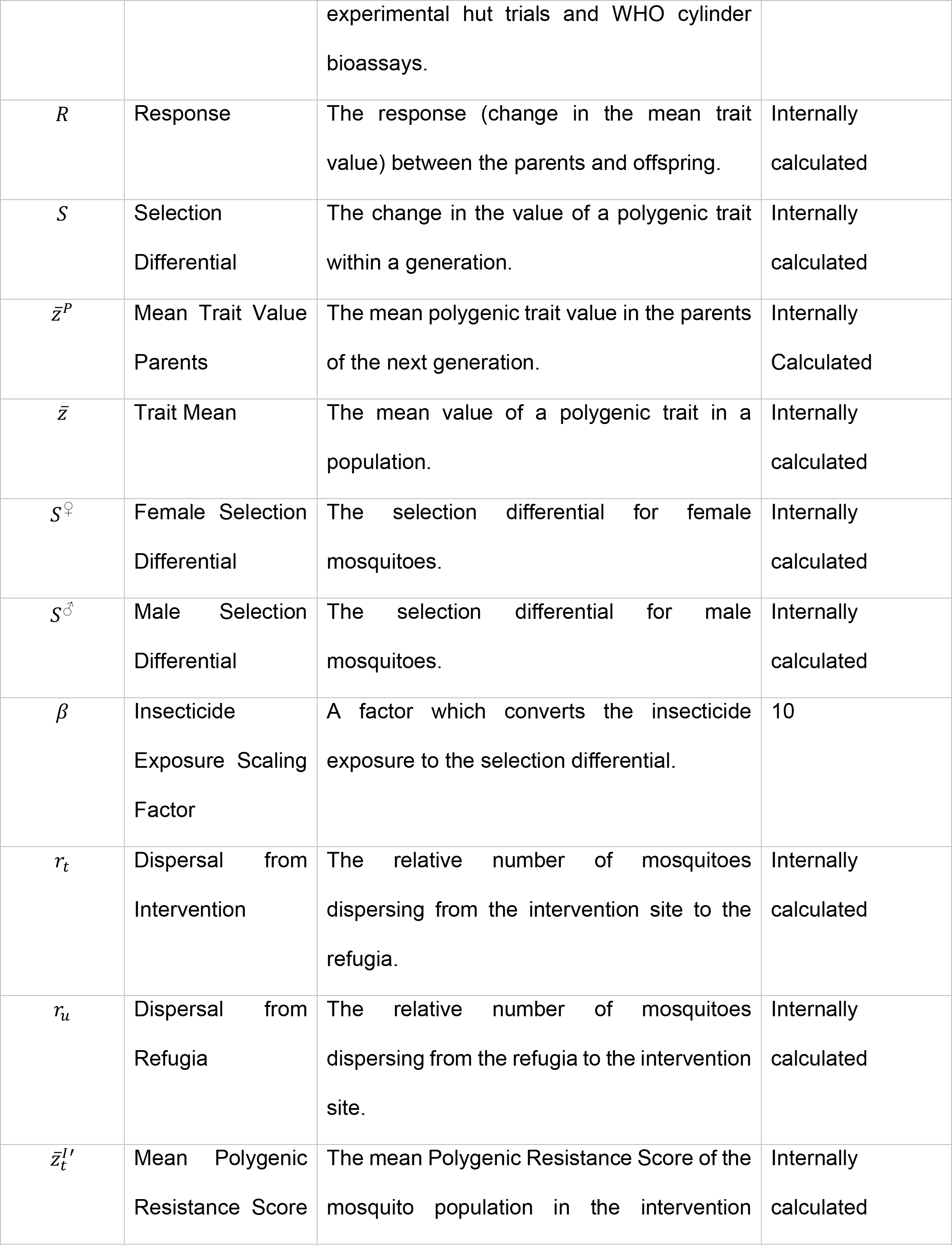
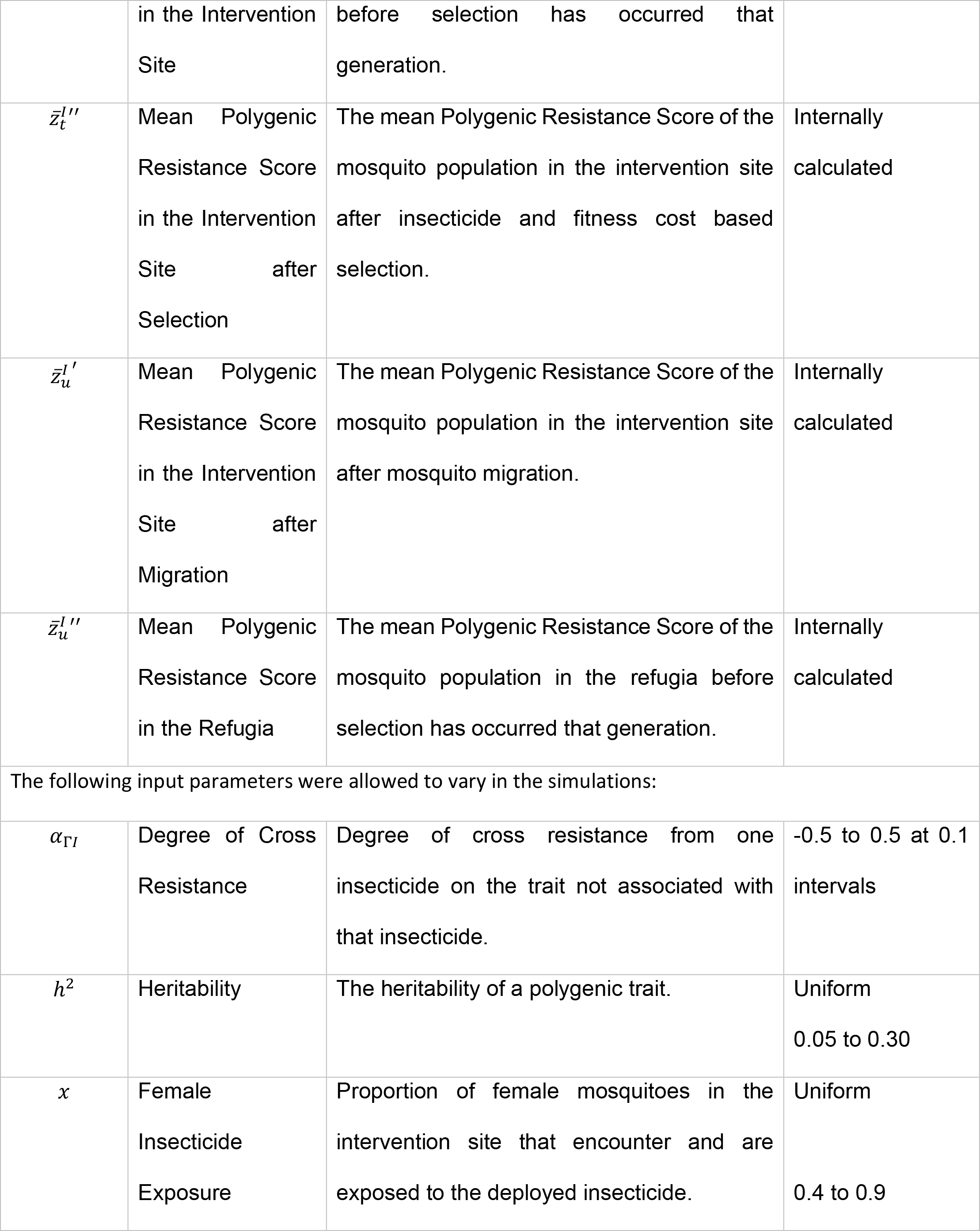
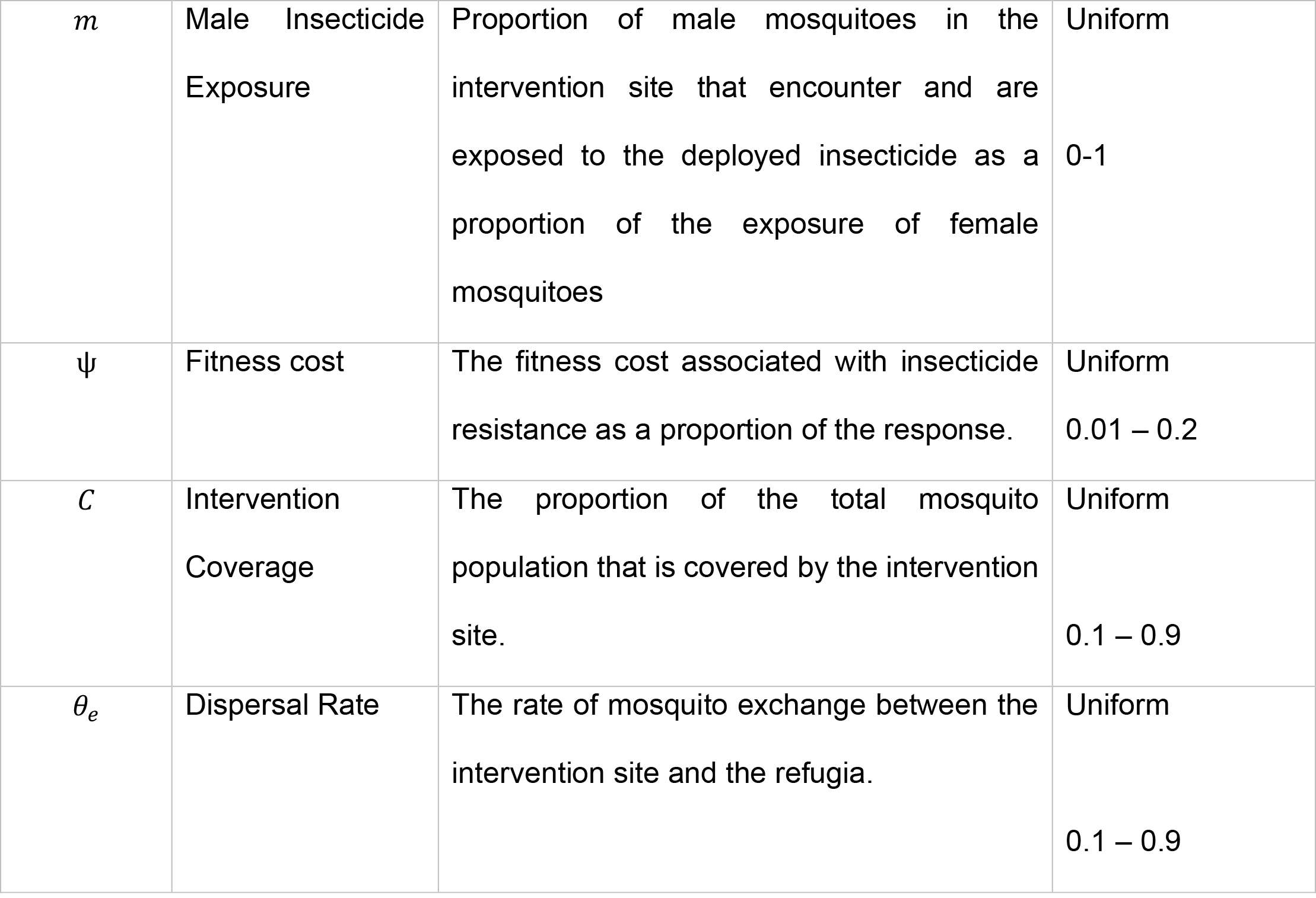
Quantitative Genetic Model Parameters and Descriptions.

| Symbol | Parameter | Description | Values |
| --- | --- | --- | --- |
| $K_i^B$ | Bioassay Survival | The probability of a mosquito with a Polygenic Resistance Unit Score of $z_I$ surviving in a WHO cylinder bioassay to insecticide $i$. | Internally calculated. |
| $K_{max}$ | Michaelis-Menten Parameter | The maximum proportion of mosquitoes that can survive in the bioassay. | 1 |
| $z_I$ | Polygenic Resistance Score | The Polygenic Resistance Score of trait $I$ to insecticide $i$. | Internally calculated |
| $z_{50}$ | Half Population Survival | The Polygenic Resistance Score which gives a 50% survival probability in a WHO cylinder bioassay. | 900 |
| $n$ | Slope of the Michaelis-Menten | The slope in the Michaelis-Menten equation | 1 |
| $K_i^F$ | Field Survival Probability | The survival probability in the field to insecticide $i$. | Internally calculated |
| $\varphi_1$ | Field-Bioassay linear model coefficient | A linear model coefficient obtained from performing a linear model on paired experimental hut trials and WHO cylinder bioassays. | 0.48 |
| $\varphi_2$ | Field-Bioassay linear model intercept | The linear model intercept obtained from performing a linear model on paired experimental hut trials and WHO cylinder bioassays. | 0.15 |
| $R$ | Response | The response (change in the mean trait value) between the parents and offspring. | Internally calculated |
| $S$ | Selection Differential | The change in the value of a polygenic trait within a generation. | Internally calculated |
| $\bar{z}^P$ | Mean Trait Value Parents | The mean polygenic trait value in the parents of the next generation. | Internally Calculated |
| $\bar{z}$ | Trait Mean | The mean value of a polygenic trait in a population. | Internally calculated |
| $s_{\text{♀}}$ | Female Selection Differential | The selection differential for female mosquitoes. | Internally calculated |
| $s_{\text{♂}}$ | Male Selection Differential | The selection differential for male mosquitoes. | Internally calculated |
| $\beta$ | Insecticide Exposure Scaling Factor | A factor which converts the insecticide exposure to the selection differential. | 10 |
| $r_t$ | Dispersal from Intervention | The relative number of mosquitoes dispersing from the intervention site to the refugia. | Internally calculated |
| $r_u$ | Dispersal from Refugia | The relative number of mosquitoes dispersing from the refugia to the intervention site. | Internally calculated |
| $\bar{z}_t^{I'}$ | Mean Polygenic Resistance Score | The mean Polygenic Resistance Score of the mosquito population in the intervention | Internally calculated |
|  | in the Intervention Site | before selection has occurred that generation. |  |
| $\bar{z}_t^{l''}$ | Mean Polygenic Resistance Score in the Intervention Site after Selection | The mean Polygenic Resistance Score of the mosquito population in the intervention site after insecticide and fitness cost based selection. | Internally calculated |
| $\bar{z}_u^{l'}$ | Mean Polygenic Resistance Score in the Intervention Site after Migration | The mean Polygenic Resistance Score of the mosquito population in the intervention site after mosquito migration. | Internally calculated |
| $\bar{z}_u^{l''}$ | Mean Polygenic Resistance Score in the Refugia | The mean Polygenic Resistance Score of the mosquito population in the refugia before selection has occurred that generation. | Internally calculated |
| The following input parameters were allowed to vary in the simulations: |  |  |  |
| $\alpha_{\Gamma I}$ | Degree of Cross Resistance | Degree of cross resistance from one insecticide on the trait not associated with that insecticide. | -0.5 to 0.5 at 0.1 intervals |
| $h^2$ | Heritability | The heritability of a polygenic trait. | Uniform<br>0.05 to 0.30 |
| $x$ | Female Insecticide Exposure | Proportion of female mosquitoes in the intervention site that encounter and are exposed to the deployed insecticide. | Uniform<br>0.4 to 0.9 |
| $m$ | Male Insecticide Exposure | Proportion of male mosquitoes in the intervention site that encounter and are exposed to the deployed insecticide as a proportion of the exposure of female mosquitoes | Uniform<br>0-1 |
| $\psi$ | Fitness cost | The fitness cost associated with insecticide resistance as a proportion of the response. | Uniform<br>0.01 – 0.2 |
| $C$ | Intervention Coverage | The proportion of the total mosquito population that is covered by the intervention site. | Uniform<br>0.1 – 0.9 |
| $\theta_e$ | Dispersal Rate | The rate of mosquito exchange between the intervention site and the refugia. | Uniform<br>0.1 – 0.9 |

### Converting Bioassay Survival to Field Survival

The model tracks the bioassay survival to insecticides via the PRS (Equation 1a). Laboratory bioassays are highly standardised using unfed, 3–5-day old female mosquitoes exposed to a fixed amount of insecticide for a fixed amount of time (WHO, 2018). This has led to concerns that bioassay results may not reflect mortality to insecticides deployed under more realistic field conditions and may have limited predictive value for operational decision-making (e,g, Grossman et al., 2020). Bioassay survival must therefore be converted to field survival to account for variable contact durations and other non-standardised factors in the field, and because insecticides select for mosquitoes capable of surviving exposure in the field and not for their ability to survive in a bioassay. Mosquito survival in bioassays has been found to be correlated with mortality in experimental huts (Churcher et al., 2016). To obtain estimates for the relationship between both as measures of survival, we therefore created a simple linear model to convert bioassay survival to survival in experimental huts which acts as our estimate for field survival to provide estimates for our mathematical model.

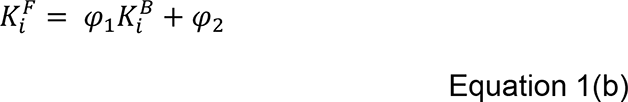

Where *K*^*F*^_*i*_ is the survival to insecticide *i* in experimental huts, which we use as an approximation for field survival. *K*^*B*^_*i*_ is the bioassay survival to insecticide *i*, (which we calculate from equation 1a in our simulations) φ_1_ = 0.48 and φ_2_ = 0.15 are regression coefficients obtained from our linear model (see Supplement 1 for details). This means a fully susceptible population (with *z̅*=0) would be expected to have an average of 15% survival to the insecticide in experimental huts. This seems intuitive as individuals vary both in their own, largely environmentally-determined, phenotypes that affect insecticide susceptibility (e.g., size, age, blood-feeding status) and the insecticidal environment they encounter i.e., concentration of insecticide (LLIN/IRS age) and duration of contact. Estimates are based on the relationship between paired female mosquito WHO cylinder bioassay survival and survival in experimental huts, and we assume this relationship also holds for male mosquitoes. We would like to highlight the lack of experimental hut studies which report male mosquitoes (i.e., numbers caught and mortality rates), which from an IRM perspective makes parameterising models challenging as we are unclear on the level of selection applied to male mosquitoes.

### The Response to Selection

The changes in population mean PRS (*z̅*) are tracked using the Breeder’s equation. The Breeder’s equation is synonymously known as the Lush equation (Walsh & Lynch, 2018):

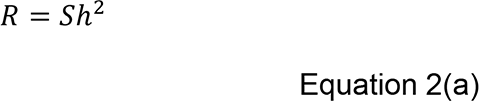

*R* is the response to selection, i.e., the inherited change in the mean value of the PRS between generations. *h*^2^is the narrow sense heritability of the trait and *S* is the selection differential. The selection differential is the within-generation change of the mean PRS of the population due to insecticide selection.

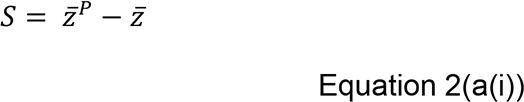

Where *z̅*^*P*^ is the mean PRS of the parents (i.e., those that either do not encounter insecticide, or encounter and survive insecticide, and go on to produce offspring), and *z̅* is the mean PRS prior to selection. Only female mosquitoes blood-feed, and because many control measures target host-seeking females it is likely that there will be sex-specific differences in insecticide exposure, and hence selection pressure. This can be incorporated by allowing the Breeder’s Equation to account for sex-specific selection (Walsh & Lynch, 2018):

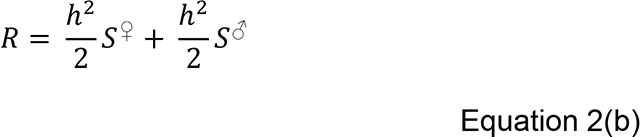

The selection differential for females and males is:

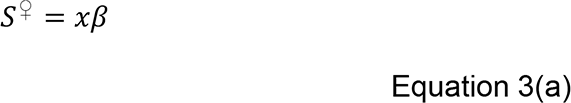

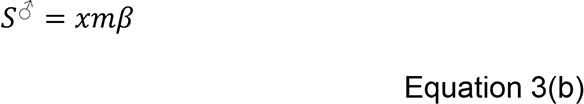

Where *x* is exposure to the insecticide (proportion of females encountering the insecticide in a generation) and beta is a scaling factor (see below). The term *m* is the proportion of male *An. gambiae* exposed to the insecticide as a proportion of the female mosquitoes, which therefore accounts for female *An. gambiae* mosquitoes being more likely to encounter insecticides especially in the form of an LLIN or IRS. The male and female selection pressures can then be implemented back into equation 2(b):

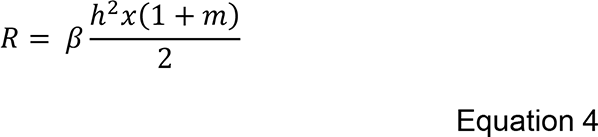

Note that insecticide deployments that do not differentially impact males and females (e.g., spraying larval breeding sites) could be investigated simply by setting *m*=1.

An important operational difficulty in applying the breeder’s equation (i.e., 2a) to selection in the field is that neither the selection differentials imposed by insecticides in the field, nor the field heritability of traits are known (heritability in the field will be much lower than in the laboratory due to the much larger environmental variance in field heritability). We therefore included an exposure scaling factor (*β*) to incorporate this uncertainty.

The value of *β* was calculated empirically so that, on average, insecticides under continuous usage would have an “insecticide lifespan” of approximately 100 mosquito generations (∼10 years) in the absence of fitness costs or migration. We define “insecticide lifespan” as the time to the withdrawal threshold of 10% bioassay survival. This was achieved by sampling the parameter space of the female insecticide exposure (*x*), male insecticide exposure (*m*) and heritability (ℎ^2^) to calculate values of the response (*R*) using equation 4b. The value of *β* was changed until this distribution centred around *R*=1, which was found to be *β*=10 (Figure 2). This is the critical part that calibrates/validates our PRS scale of z. It means that if z=0 at the start of the simulation (i.e., IR is absent) it should usually reach a value of z=100 (giving 10% bioassay survival, see Equation 1(a)) after around 10 years of continuous deployment. The value of 10 years was selected as a reasonable time, based on experience (and expectation) that insecticides take roughly 10 years (i.e., ∼100 generations) of deployment before starting to fail (a >10% survival rate is regarded as indicative of potential for future failure (WHO, 2018)). It is, however, user-defined, and the simulations could be recalibrated by changing *β* so that the “insecticide lifespan” is reached (i.e., 10% bioassay survival) most frequently occurs after 5, 15, 20 years of deployment according to operator beliefs on likely timescales and the threshold at which insecticide withdrawal should occur.

**Figure 2.**
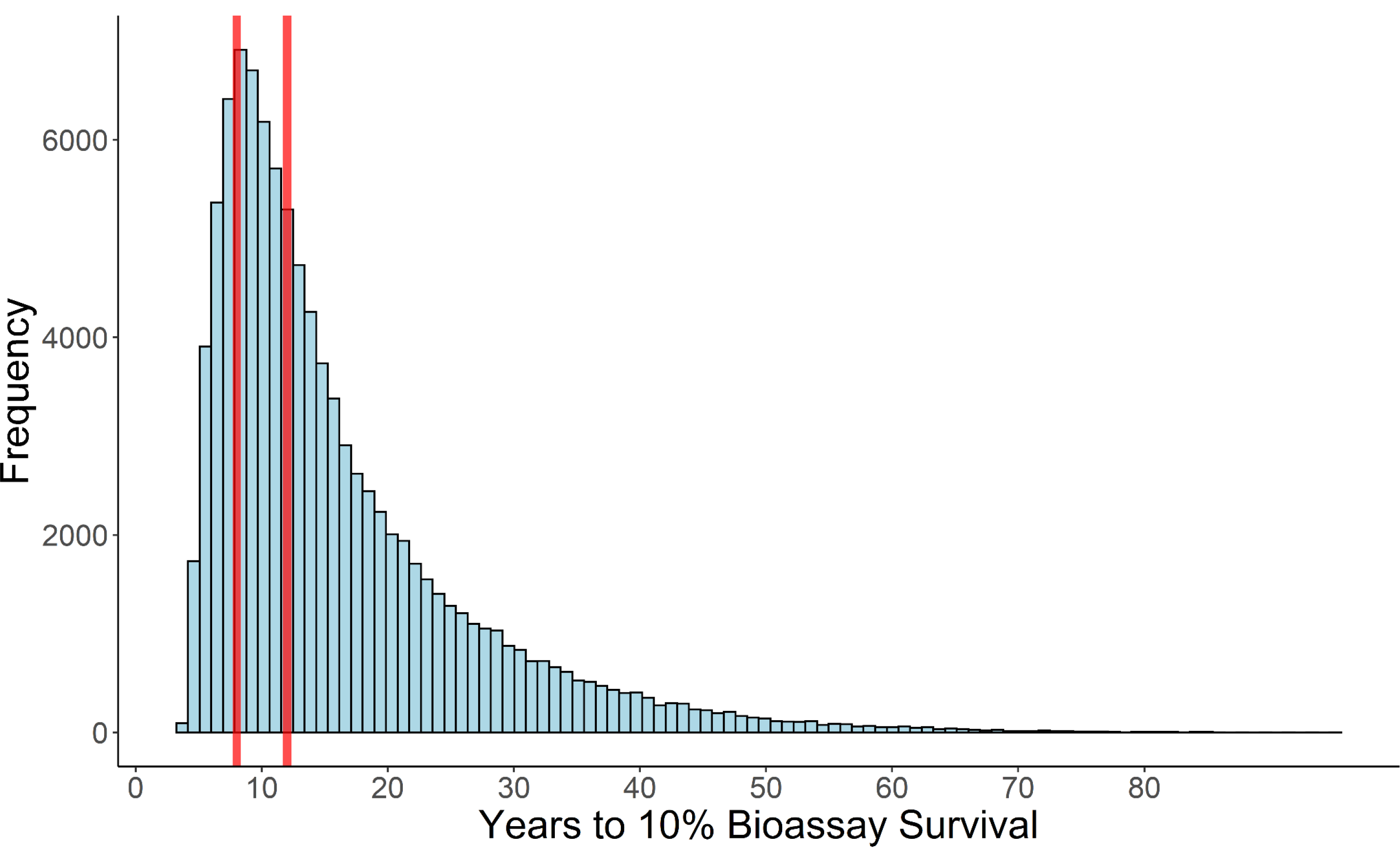
Histogram of the Insecticide Lifespan Assuming the Exposure Scaling Factor (β) = 10. Calibration of the exposure scaling factor is needed to convert selection to the desired timescale of expected insecticide lifespans (i.e., Equation 4B). The exposure scaling factor was varied until the peak of the histogram was approximately 10 years, as we expect insecticides to last this length. The red vertical lines are at 8 and 12 years, indicating the range inside which we would expect insecticides to most frequently reached the reach the 10% bioassay survival withdrawal threshold and therefore reach the end of its insecticide lifespan.

### Fitness costs

Insecticide resistant mosquitoes may be less fit than insecticide susceptible individuals (Osoro et al., 2021; Tchouakui et al., 2020). The simplest way of incorporating a fitness cost would be to set it as an absolute value defined as the reduction in *z* to be applied each generation. However, the same absolute decrease in *z* could be sufficient to prevent the evolution of resistance in simulations where selection is weak or could have only a marginal impact in simulations where selection is very strong. We therefore incorporated a fitness cost by defining it proportional to response in the simulation (see Equation 4b) i.e.

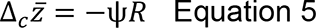

Importantly any given value of ψ now has the same impact in all runs when simulating over numerous parameter combinations, making its effect constant and hence its impact can be better estimated. Δ_*c*_*z̅* is the change in the mean PRS per generation as a result of the fitness costs associated with the resistance genes (ψ), noting that is trivial to set Δ_*c*_*z̅* = 0 if costs are assumed to be absent. We first calculate the response as though the insecticide was present (in the intervention site), before multiplying this response value by the fitness costs. This helps to ensure the fitness cost is smaller than the response allowing resistance to take-off when *z* is small. It is assumed that the fitness costs remain constant regardless of the magnitude of the IR level, *z*. This is because fitness costs arise from competition between individuals within the Normal distribution of the population. Since we assume the standard deviation of the Normal distribution does not change with the magnitude of the population mean *z̅*, the level of competition will be constant, and the fitness costs are independent of IR (i.e., competition is not between resistant individuals and fully susceptible individuals as occurs when assuming IR is a monogenic trait). Note that we assume that *z̅*=0 is evolutionary stable (i.e., resistance levels are at their basal, pre-deployment level) and fitness costs cannot reduce *z̅* to less than zero, for example when the insecticide is not being deployed.

### Accounting for Mosquito Dispersal

Mosquitoes may disperse between the intervention site and an insecticide-free refugia. Dispersal from intervention to refugia is given in Equation 6a and from refugia to intervention site Equation 6b (see Supplementary Information in Hastings et al., 2022 for details).

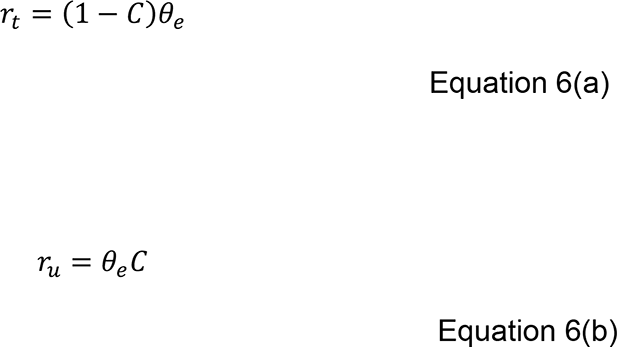

Where *r*_*t*_ is the number of mosquitoes migrating from the insecticide treated location to the insecticide-free refugia. and *r*_*u*_ is the number of mosquitoes migrating from the insecticide-free refugia to the insecticide treated location. *C* is the coverage of the insecticide (i.e., the proportion of the population covered by the intervention), and *θ*_*e*_ is the proportion of mosquitoes dispersing.

### Tracking the Polygenic Resistance Score

The impact of the insecticide selection pressure and fitness costs associated with IR during insecticide deployment in the location where the insecticide is deployed is quantified as the change over a generation and is given by the response to insecticide selection and the fitness costs of IR as explained previously.

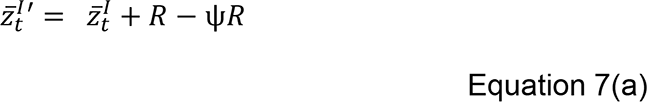

Where the superscript *I* represents the insecticide-resistance trait (resistance to insecticide *i*), and the subscript *t* represents the intervention site.

After the effect of insecticide selection and fitness costs are implemented, dispersal occurs:

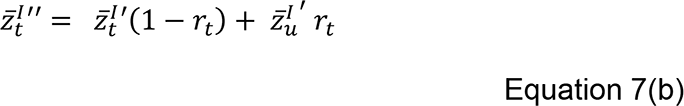

The mean PRS in the site depends on the PRS in the proportion of the mosquitoes staying in the treated site *z̅*^*I*^′(1 − *r*_*t*_) and the proportion of the mosquitoes that have immigrated from the refugia to the insecticide treated location. 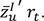.

When an insecticide is not deployed in the treatment location (e.g., because it has been rotated out) there is no insecticide selection but there are fitness costs which are calculated as explained previously.

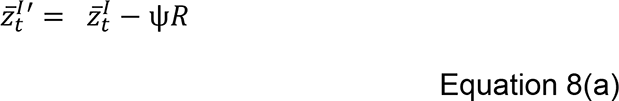

The impact of mosquito dispersal is then allowed as in equation 7(b) i.e.:

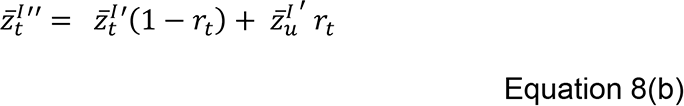

In the insecticide-free refugia (subscript *u*), only IR costs are present as there is never any insecticide deployment.

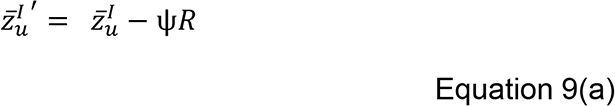

Migration is also allowed to occur, with mosquitoes dispersing from or to the refugia.

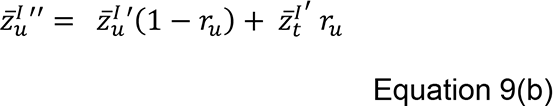

Where 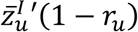 is the mean resistance of the individuals staying in the refugia site and 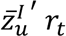 is the mean resistance of those joining from the intervention site.

Note the use of primes: the single prime (e.g., *z*^*−I*^ ^′^) indicates the value of z after insecticide selection and the double prime (e.g., *z*^*−I*^ ′′) is the value after both insecticide selection and mosquito dispersal. It is the double prime value that becomes the mean PRS value in the next generation.

### Special Case 1: Cross resistance and cross-selection between insecticides

Cross resistance between insecticides is often considered an important factor when selecting insecticides for an IRM strategy especially relevant to the mixture strategy (e.g. Curtis, 1985), yet is often not included in mathematical models evaluating IRM strategies (Rex Consortium, 2010). Via (1986) included genetic correlation, but in a model which tracked the median lethal dose (LD50) to the respective insecticides and therefore needed a more complex method to calculate the genetic correlation.

We note that cross selection would refer to the amount of selection one insecticide would cause for resistance to another. Whereas cross resistance is the amount of resistance a trait for one insecticide would confer to another insecticide. These two phrases (cross selection and cross resistance) are often interchanged; but to aid our readers we use the term cross resistance as we expect readers to be more familiar with this concept.

The degree of genetic correlation between the level of IR to insecticide *i* (trait *I*) and *γ* (trait Γ) is quantified as (α_*I*Γ_). As all the traits measured in this model are on the same scale (e.g., *z*=100 for trait *I* is 10% bioassay survival to insecticide *i*, and z=100 for trait *J* is 10% bioassay survival to insecticide *j*), there is no need for regression/variance coefficients to translate between different scales as would arise if, for example, finding the degree of cross selection between selection of wing length (mm) and body mass (mg) as the two traits are on different scales), and we can assume a simple correlation between the traits. For example, if the genetic correlation is 10% between trait *I* and trait *J*, then insecticide selection causing a one-unit increase in *z̄* would cause a 0.1 unit increase in *z*_*J*_.

Equation 8(a) is updated to include cross resistance/selection, such that when insecticide *i* is not deployed and insecticide *γ* is deployed, there is also selection on trait *I* from insecticide *γ* based on the genetic correlation trait Γ and trait *I*, termed *α*_ΓI_.

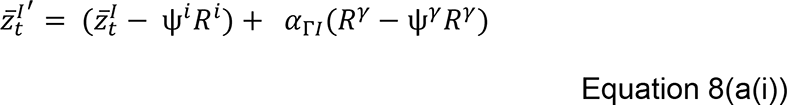

Where *γ* ∈ {*j*, *k*, *l* … }, the set of the other insecticides that may be deployed in the mixture.

Where Γ ∈ {*J*, *K*, *L* … }, the set of traits that correspond to IR to the corresponding insecticide.

Equation 8(a(i)) is passed to equation 9(a) as previously described to allow migration to occur as previously described. The refugia equations do not need updating as we assume no insecticides under consideration in the IRM are deployed in the refugia.

### Special Case 2: Inclusion of Mixtures

Insecticide mixtures describe the simultaneous deployment of two (or, in theory, more) insecticides in the same formulation. Given that LLINs and IRS formulations currently submitted for WHO pre-qualification do not include more than two insecticides, mixtures are limited to contain only two insecticides in our simulations. The primary idea behind mixtures is that if one insecticide in the mixture fails to kill the mosquito, the other insecticide of the mixture will do so. This requires equation 7a to be updated. When an insecticide mixture contains both insecticides *i* and *j* the mosquito must survive the encounter with one part of the insecticide before selection can occur. In this example, prior to selection on trait *I* the mosquito must first survive its encounter with insecticide *j*.

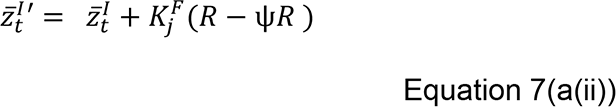

Where *K*^*F*^_*j*_ is the average probability of a mosquito surviving exposure to insecticide *j* in the field when deployed at the full dose, which depends on the concurrent value of *z*_*J*_ and is obtained from equation 1b. When insecticide *i* is not deployed in a mixture, then the model proceeds through equation 8a as normal. The refugia equations 9(a) and 9(b) remained unchanged apart from being passed the updated *z̅*^*I*^′ from equation 7(a(ii)).

### Special Case 3: Cross resistance and Selection with Mixtures

When an insecticide mixture is deployed that consists of insecticide *i* and insecticide *j* and there is cross resistance between insecticides there is both direct and indirect selection on trait *I* which is then scaled by the survival probability to the second insecticide in the mixture.

Equation 7(a(ii)) is therefore updated so that the survival probability to the second insecticide in the mixture acts on the changes in resistance intensity.

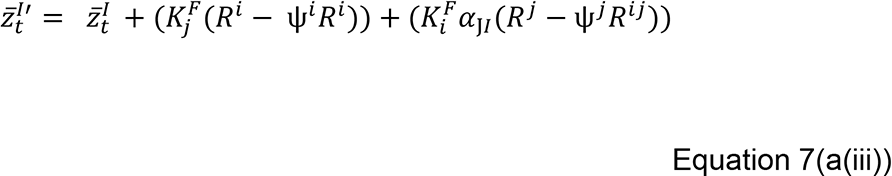

Where the second term incorporates direct selection, and the third term incorporates indirect selection by cross resistance. Note that if cross-selection is absent *α*_ΓI_ =0 so equation 7a(iii) is, as expected, equivalent to Equation 7a(ii).

Our methodology can be extended to track more than two insecticides so it would technically be possible for the tracked insecticide to not be deployed in the currently deployed mixture. This would occur if insecticide *i* is being tracked, while a mixture containing insecticide *j* and *k* was deployed. In this instance there is indirect selection on trait *I* from insecticide *j* and insecticide *k*. There is therefore indirect selection on trait *I* from insecticide *j* scaled by survival to insecticide *k*. There is also indirect selection on trait *I* from insecticide *k* scaled by survival to insecticide *j*.

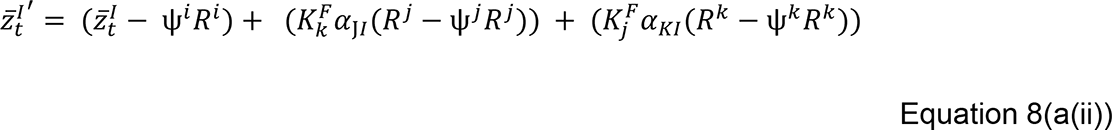

Note, in our simulations, we only use two insecticides, therefore all mixture simulations use only equation 7a(iii), however equation 8a(ii) is presented for completeness and demonstrates this modelling framework can extend to allow simulations with more than two insecticides.

These are then passed to equations 7(b) and 8(b) to allow for migration. The refugia equations 9(a) and 9(b), remain unchanged besides being passed the updated *z̅*^*I*′^from either equation 7a(iii) or 8a(ii).

The mathematical model described above is coded in R (R Core Team, 2020), version

4.0.3. Model code is written following modern coding practices including the use of modular coding, unit testing and maintenance in a version-controlled repository.

### Description of the Simulations

We define several key terms and model rules here:

*Insecticide Armoury:* The number of different insecticide formulations available for deployment. Only insecticide formulations in the armoury can be deployed. Insecticides can be withdrawn from the armoury and returned to the armoury based on the withdrawal threshold and return threshold described below.

*Withdrawal Threshold*: Is the resistance threshold whereupon an insecticide is considered to have failed and is withdrawn from the armoury. We set the withdrawal threshold at 10% bioassay survival (90% bioassay mortality) as a ≥10% bioassay survival indicates there is confirmed resistance in the mosquito population (WHO, 2018). The withdrawal threshold can be set by the user to be higher or lower.

*Return Threshold*: The bioassay survival an insecticide must reach for a previously failed insecticide to be return to the insecticide armoury and become re-available for deployment. This is to prevent an insecticide that has recently failed being immediately returned to deployment. In our presented simulations the return threshold was set at 8% bioassay survival, though this can be set by the user. For example, if an insecticide reaches the withdrawal threshold of 10% bioassay survival the insecticide remains unavailable for re-deployment until its bioassay survival falls below 8%. We use 8% as this value allows withdrawn insecticides to have the potential redeployed, as lower return thresholds are unlikely to ever be reached in the simulations. The withdrawal threshold value can therefore impact the difference between sequences and rotations, if set too low a withdrawn insecticide (in sequence) is not returned to the armoury before the second insecticide is withdrawn. An illustrative example of the process of withdrawing and returning insecticides is presented in the sequence simulation example in Figure 3.

**Figure 3.**
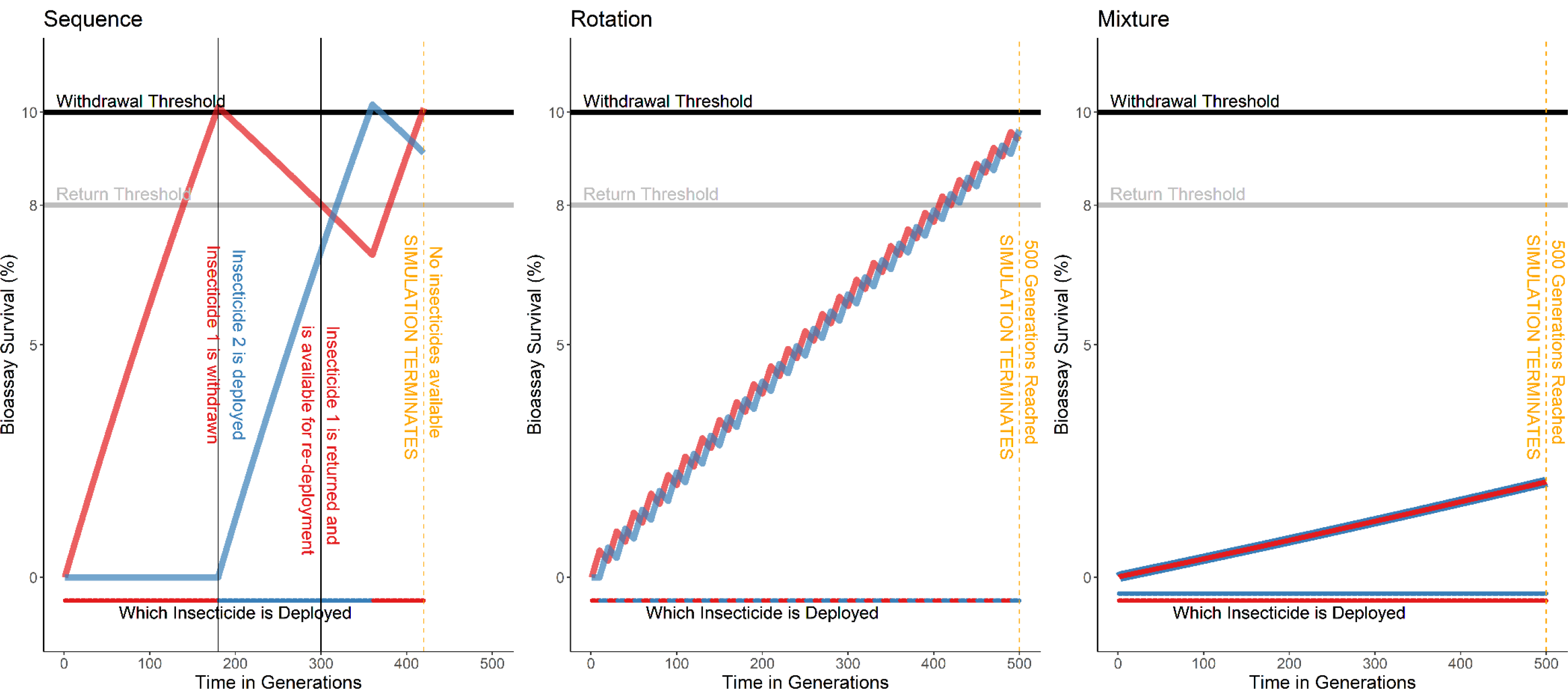
Example Simulations of the Sequence, Rotation and Mixture Strategies. ***Sequence strategy:*** Insecticide 1 (red) is deployed continuously until it reaches the withdrawal threshold and is replaced by the Insecticide 2 (blue) at the next deployment opportunity. Insecticide 2 is deployed continuously until it too reaches the withdrawal threshold. When Insecticide 1 is not deployed, fitness costs reduce resistance. Once the resistance has reached the return threshold (at 300 generations), it becomes re-available for deployment. The sequence simulation terminates at 420 generations as neither insecticide is available for deployment. ***Rotation Strategy:*** The deployed insecticide deployed is changed at each deployment interval. The simulation was terminated at 500 generations, with both insecticides still available for deployment as neither had yet reached the withdrawal threshold. ***Mixture Strategy:*** Insecticide 1 and 2 are deployed together as a single formulation, in a de facto sequence. The simulation was terminated at 500 generations, with both insecticides still available for deployment as neither had yet reached the withdrawal threshold. These simulations are presented only to explain how each IRM strategy works. The parameter inputs were: deployment interval = 10 generations, cross resistance = 0, start resistance = 0, heritability = 0.2028686, male insecticide exposure = 0.2500806, female insecticide exposure = 0.8645515, fitness cost = 0.1730878, intervention coverage = 0.6885651, dispersal = 0.8844445.

*Deployment Interval / opportunity:* The deployment interval is the timeframe (in mosquito generations) between insecticide deployments and deployment decisions. For LLINs this is 30 generations (∼3 years) and for IRS 10 generations (∼yearly), hence there is only a deployment opportunity (i.e., the opportunity to change the deployed insecticide) every 3 or 1 year. The model does allow for a user input of the deployment interval, allowing for 2 IRS sprays a year (for locations with two transmission seasons) or more frequent replenishment of LLINs. The deployment interval is constant throughout each simulation.

IRM Strategies:

The three primary IRM strategies evaluated are sequences, rotations, and mixtures. Illustrative examples for each IRM strategy are given in Figure 3.

- *Sequence*s: the first insecticide is deployed continually until it reaches the withdrawal threshold. At which point it is withdrawn at the next deployment opportunity and replaced with the second insecticide. The second insecticide is then deployed continually until it also reaches the withdrawal threshold. If the first insecticide has fallen below the return threshold of 8% WHO bioassay survival it can be redeployed.
- *Rotations*: the insecticide in deployment is switched at each deployment interval. In this strategy, the simulation also stops when the insecticide is unable to be rotated out and replaced, that is if previously deployed insecticide would be deployed in sequence. This deployment restriction is relaxed for simulations where unique insecticides are used (see Adaptive Rotations description later).
- *Mixtures*: both insecticides are deployed simultaneously within a single formulation and the mixture is therefore deployed as a sequence. The mixture consists of both insecticides being deployed at their recommended single-formulation dosage. The simulation stops either when bioassay survival to either insecticide in the mixture exceeds 10% (even if overall mortality to the mixture remains below the withdrawal threshold), or at 500 generations. We do this for two main reasons: (i). Diagnostic doses have thus far been made only for the single insecticide, rather than the mixture. (ii). Procurement/logistical failures leading to the deployment of just a solo insecticide, and therefore the insecticide is still available.

For simulations where insecticides are given unique properties, we must adapt the rotations strategy so that this strategy does not appear artificially ineffective due to the previously described rotations rules:

- *Adaptive rotations:* In adaptative rotations we relax the restriction that prevents an insecticide being immediately redeployed. This means when only one insecticide is available for deployment (because the other is above the return threshold), it continues to be deployed. However, at the first deployment opportunity where the other insecticide becomes available (i.e., has fallen below the return threshold), the strategy defaults back to rotations.

We limit our simulations to containing only two insecticides as this is the current number in mixture IRS and LLINs. We therefore look at purely evaluating the differences between strategies, and not the complexities associated with allowing for additional insecticides and extending the armoury of strategies. We note that the research, development, and economic complexities with creating a mixture formulation may not be equivalent to creating two separate single insecticide formulations.

### Designing and Running Simulations to Directly Compare Strategies

This set of simulations assumes each insecticide in the parameter combination has the same properties (i.e., heritability, fitness cost and starting resistance are the same for both insecticides). This allows for better comparison of the effect of the IRM strategy itself as the efficacy is not dominated by a single insecticide to which resistance evolves very slowly or inhibited by an insecticide to which resistance evolves very rapidly (this assumption is relaxed in the next section).

The parameter space is described on Table 2 and was sampled using Latin hyperspace sampling (Carnell, 2020). A total of 5,000 randomly generated parameter inputs were used. Cross resistance/selection values were used from -0.5 to 0.5 at 0.1 intervals, giving 11 different values. Simulations were started assuming both insecticides were novel (z=0, in both intervention site and refugia) or both insecticides had previously been used and there was some resistance present (z=50, corresponding to ∼5% bioassay survival. Therefore, a total of 220,000 (5,000 random parameter values x 11 cross resistance values x 2 starting resistance values x 2 deployment intervals) unique parameter inputs were available for each IRM strategy. The same 220,000 parameter inputs were used for each IRM strategy to allow for direct comparisons between strategies. Three IRM strategies were evaluated so a total of 660,000 (220,000 x 3) simulations were conducted.

### Simulations with Unique Insecticide Properties

We ran a set of simulations which allowed for each insecticide in the armoury to have different, unique properties. In these simulations the two insecticides were allowed to have different heritability, fitness cost and starting resistance (PRS). For the starting PRS, this was sampled within a uniform distribution between 0 and 80 (ensuring the PRS was below 100 as this is our criteria for a failed insecticide). For each insecticide, the starting PRS was the same in both the intervention site and the refugia. Cross resistance was included between the insecticides and was sampled within a uniform distribution between -0.5 and 0.5. Latin hyperspace sampling was used again (Carnell, 2020). This allows for the comparison of IRM strategies under more realistic conditions, where insecticides have different properties and may have been subject to previous selection and have different rates of evolution. A total of 50,000 simulations were run for each of sequences, rotations, mixtures and the adaptative rotation strategy. The same 50,000 parameter sets were used for each strategy, allowing for direct comparisons between the strategies. Therefore 200,000 (50,000 parameter sets x 4 IRM strategies) simulations were run.

### Outcome Measures

One challenge with any study of different strategies/interventions/policies is defining how best to evaluate them (see discussions in Madgwick & Kanitz, 2022a). For evaluating IRM we consider that there are three potential main outcomes of interest. First is the “strategy lifespan”, second is the average level of resistance and third is the highest level of resistance achieved. We consider the “strategy lifespan” to be the primary outcome.

*Strategy Lifespan*: The total duration of the simulation in generations. The simulation stops either when (i). neither insecticide is available for deployment (because both are above the return threshold) or (ii). when the simulation has run for 500 generations. The reason for capping at 500 generations is if there are no obvious difference between deployment strategies at this point, which represents a ∼50-year time horizon, then the strategies are equivalent over any notional policy timeframe. When one strategy has a longer strategy lifespan than another for the same parameter inputs, we define that strategy as having won and the other strategy to have lost. If two (or more) strategies have the same strategy lifespans, we say they have drawn. We presume the longer an IRM strategy lasts, the better the IRM strategy. We report the number of wins, losses and draws as percentages of the totals. We also note the size of the win in terms of % difference in lifespan on the insecticides.

We expect a strategy to have a strategy lifespan of at least 10% longer to justify the using a potential more logistically complex or economically expensive IRM strategy. We therefore define any win where the strategy lifespan increased by 10% or more to another as an “operationally relevant win”. This is especially the case where the benefit of one IRM strategy over another may not be seen for over 30 years, which is beyond the timeframe used for operational planning. This threshold was used as a soft cut-off as used similarly by Madgwick & Kantiz (2022).

If the strategy lifespan is ≥10% longer than another strategy (under the same conditions) we describe that strategy as having an “operationally relevant win”. This helps to identify simulations where there would likely be a benefit in choosing one strategy over another.

When two strategies have equal strategy lifespans (for the same parameter inputs), we say they have drawn. Draws can occur if both simulation runs out of insecticides (i.e., all insecticides have bioassays survival <10%) at the same timepoint, or a pair of simulations both reach the 500-generation maximum. Where the sets of simulations draw, the secondary outcomes can be compared. Secondary outcomes are compared only in simulations with equal strategy lifespans.

The secondary outcomes are:

1. The mean bioassay survival to the currently deployed insecticide(s) during the simulation. We report this rather than the corresponding field survival (Equation 1b) as bioassay survival is measured in operational settings to inform decisions. We assume insecticide-based interventions are more effective when bioassay survival is lower as the killing effect on the population size and age structure would be greater. Therefore, a lower mean bioassay survival is preferrable.
2. The peak bioassay survival reached during the simulation by any insecticide at any timepoint. As bioassay survival increases, we may expect an increase in the potential for compensatory genes to evolve to counteract the effect of fitness costs, and therefore a lower peak bioassay survival is preferable.

Following Levick et al (2017), we stress that modelled estimates for the longevity and performance of strategies are not absolute predictions of how long a particular deployment would last in practice, instead serving as a general guide as to which strategy would be expected to perform better than another under such conditions.

### Statistical Analysis

All statistical analysis and data visualisation was conducted in R version 4.0.3 (R Core Team, 2020). The following R packages were used: epiR (Stevenson et al., 2020) for partial rank correlation, mgcv (Wood, 2011) for generalised additive models, MASS (Venables & Ripley, 2002) for the negative binomial GLM, and randomForest (Liaw & Wiener, 2002) for the random forest models. Data visualisation used ggplot2 (Wickham, 2009). We separate our analysis into three parts.

- First, we directly compare strategies deploying a single insecticide at a time (sequences and rotations) against one another.
- Second, we compare deploying insecticides singularly (sequences and rotations) against deploying the same insecticides together in a mixture. This is done because not all insecticides may be chemically suitable to form mixtures with all other insecticides and because deployment as mixtures may substantially increase deployment costs.
- Thirdly, model sensitivity analysis was conducted using partial rank correlation, generalised linear modelling, and random forest models to identify parameters driving IR and to identify parameters which may offer predictive value in informing the IRM decision-making process.

### Analysis to Identify Parameters Driving Insecticide Resistance and to Inform Decision Making

#### Partial Rank Correlation

Partial rank correlation was used to assess the degree of correlation between the randomly generated parameter input values and the strategy lifespans of the IRM strategies where the insecticides were given equal properties. This identified the main factors driving the evolution of IR and was conducted separately for each IRM strategy, each level of starting resistance, and each cross-selection value. Two-sided partial rank correlation was used to determine the direction of the correlation. Correlation estimates were calculated with a 95% confidence interval.

#### Generalised Linear Modelling and Generalised Additive Modelling

Simulations which ran to completion (i.e., terminated at 500 generations) were excluded from the analysis as these simulations were artificially terminated and their inclusion could lead to an underestimation of effect sizes. A total of 86566 simulations were included after these runs were excluded. Initial exploration with a Poisson GLM, indicated the data was over-dispersed (dispersion = 5.937677, p < 2.2e-16). A negative binomial generalised linear model was therefore fitted. The model parameters of heritability, female insecticide exposure, male insecticide exposure, intervention coverage, fitness cost and dispersal were used as predictors. The IRM strategy was input as a factor. The starting resistance was converted to a factor, with 0 being defined as a “novel insecticide” and 50 being a “pre-used insecticide”, in this set of simulations insecticides only started at either 0 or 50 PRS. The deployment interval was input as a factor. Generalised additive models were conducted to assess for non-linear relationships. Where notable non-linear relationships were found, splines were included in the model. The knot position of the splines was found through maximising the log-likelihood. Knot positions were included to a resolution of 3 decimal places.

#### Random Forest Models

Random forest models involve fitting multiple regression classification tree models, with the final model being a composite of all these individual trees. This has the benefit of no single tree being used to inform the analysis, and therefore is less impacted by the fitting of a single tree and splits occurring from the model fitting methodology of a single tree. A random forest model can then be used to identify which individual parameters provide information which gives the best predictive accuracy.

Eight random forest models were fit to predict the operational outcome. Such that the models would be predicting the optimal IRM strategy using the following comparisons.

1. Equivalent Insecticides, Sequence vs Rotations, all intervention coverages.
2. Unique Insecticides, Sequences vs Adaptive Rotations, all intervention coverages.
3. Equivalent Insecticides, Sequences vs Rotations vs Mixtures, all intervention coverages.
4. Unique Insecticides, Sequences vs Adaptive Rotations vs Mixtures, all coverages.
5. Equivalent Insecticides, Sequence vs Rotations, intervention coverage ≥0.5.
6. Unique Insecticides, Sequences vs Adaptive Rotations, intervention coverage ≥0.5.
7. Equivalent Insecticides, Sequences vs Rotations vs Mixtures, intervention coverage ≥0.5.
8. Unique Insecticides, Sequences vs Adaptive Rotations vs Mixtures, intervention coverage ≥0.5.

The models were fitted against a random sample of 70% of the respective simulations (a training dataset), and the random forest models fit to this data was used to predict the operational outcome for the remaining 30% of the samples to estimate the accuracy of the model in predicting the correct outcome. The variable importance of each of the parameters from each model is then reported and corresponds to how much the prediction error is affected by removing the parameter (Liaw & Wiener, 2002). Therefore, parameters with greater importance correspond with higher model accuracy, and are likely to be more beneficial to measure in the field as a basis for IRM decision-making. This model fitting process was repeated but restricted to include only simulations where the intervention coverage was ≥0.5. This was done to force the models to choose a strategy because, when coverage is >0.5, the no operational win outcome was no longer the dominant outcome (see later discussion of Figure 12).

## Results

### Comparing Sequences versus Rotations

When comparing the strategy lifespans at the “global level” (i.e., crude averages, not accounting for any other parameters), then the sequence and rotation strategies appear to perform broadly equally, with a similar number of wins and operational wins for each strategy (Table 3). However, when accounting for the starting resistance, deployment interval and the degree of cross resistance more distinct patterns can be observed. When the cross resistance between the two insecticides is positive, sequences become the preferable strategy to maximise the strategy lifespan. If cross resistance is zero or negative (i.e., resistance to one enhances susceptibility to the other) rotations become the favoured strategy, unless the insecticides have pre-existing resistance. In this case there is little difference with a 10-generation deployment interval, but sequences are favoured with a deployment interval of thirty generations (Figure 4). Due to the large number of draws in strategy lifespans (72% of comparisons are tied) it is useful to also consider the secondary bioassay survival outcomes. When sequences and rotations draw, rotations outperform sequences by having both lower peak bioassay survival and a lower mean bioassay survival to the deployed insecticide, regardless of the starting resistance, deployment interval, or degree of cross resistance (Figure 5). This is expected because most draws occur when both strategies last for the maximum of 500 generations and sequences by model rule definition explicitly run insecticides up to the withdrawal threshold. In contrast, rotations often hold both insecticides at lower polygenic resistance score (PRS) over the duration of the simulations.

**Figure 4.**
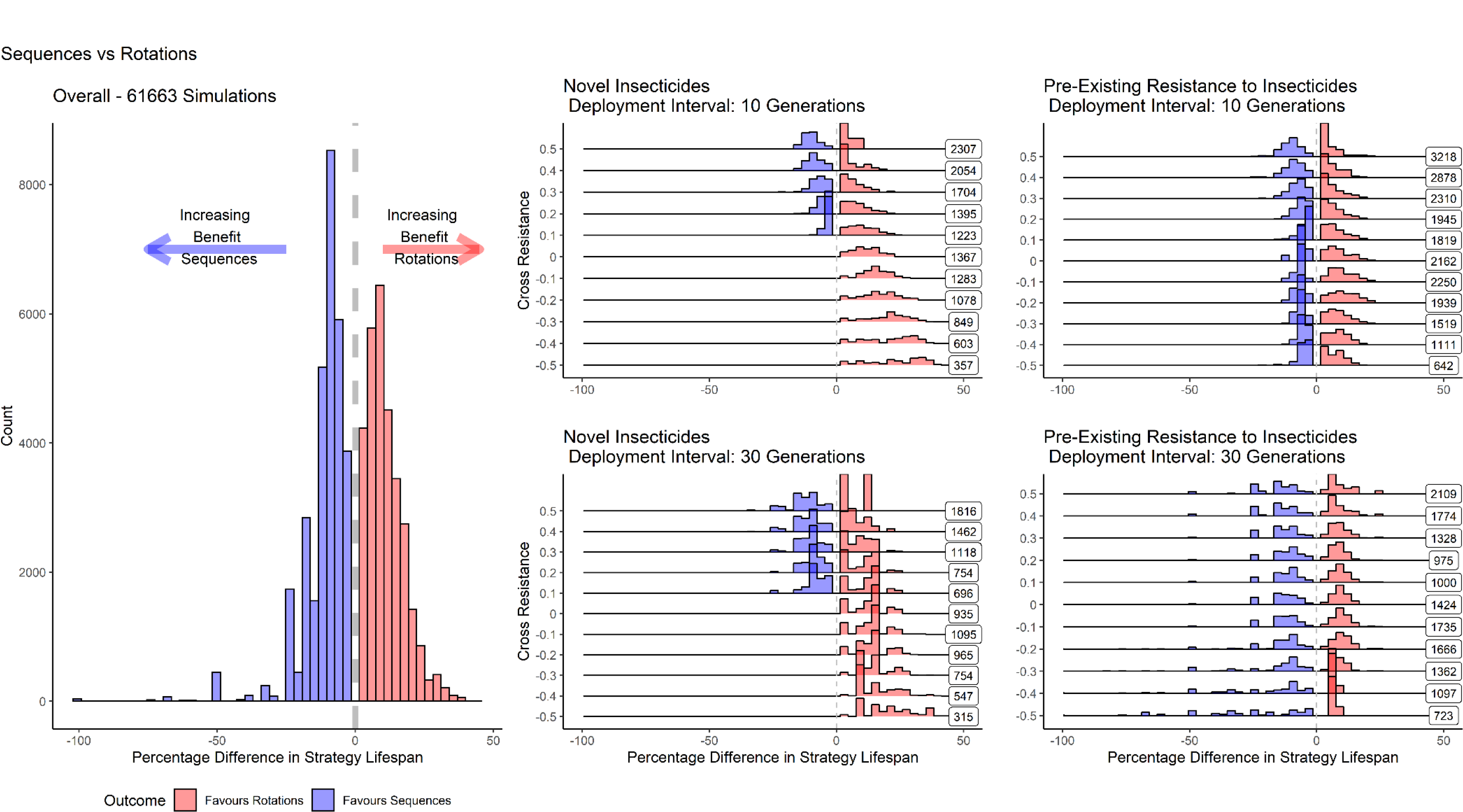
Frequency Distributions Comparing the Strategy Lifespans Between Sequences and Rotations for Insecticides with Identical Properties. Values above zero indicate rotations (red) are the favoured IRM strategy, values below zero indicates sequences (blue) were the better performing strategy. Draws (identical strategy lifespans, where the difference was 0 generations) were excluded from the plots. Left plot is the overall global distribution without any stratification. The four plots on the right are stratified by the deployment interval (top-bottom: 10 or 30 generations) and starting resistance (left-right; novel (z=0, 0% bioassay survival) or pre-used (z= 50, ∼5% bioassay survival)). Each row in these plots is the frequency distribution of the percentage difference in strategy lifespan for depending on the amount of cross resistance. For the cross-selection plots, the numbers on the right-hand side of each plot indicate how many comparisons are plotted in the distribution (that is, the number of comparisons where the simulations did not draw).

**Figure 5.**
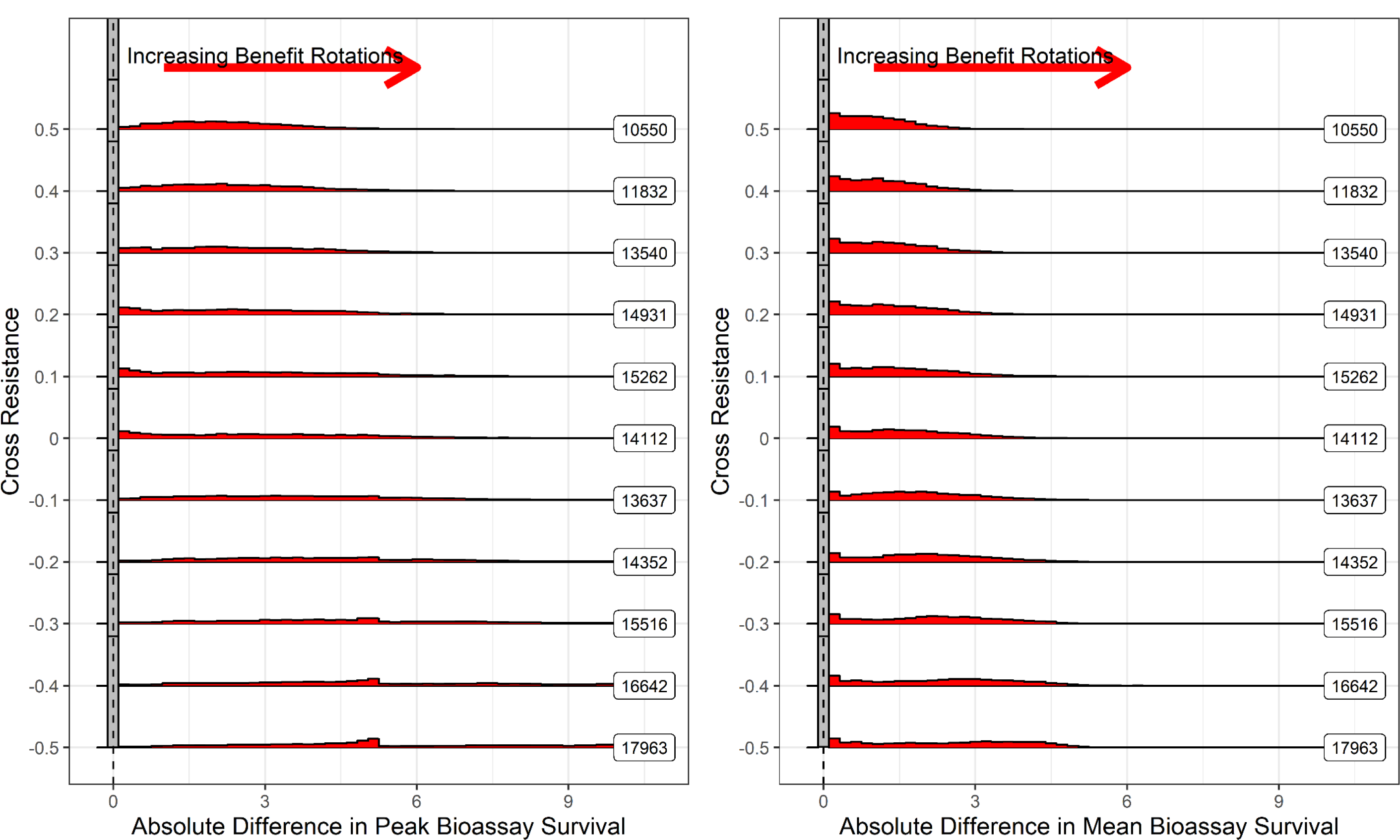
Difference in the Peak and Mean Bioassay Survival for Drawn Sequences versus Rotations Simulations for Insecticides with Identical Properties. Left Panel is difference in the peak bioassay survival, where values above zero favour rotations and values below zero favour sequences. Right panel is difference in the mean bioassay mortality to the deployed insecticide, where values above zero favour rotations and values below zero favour sequences A total of 158337 simulation pairs (same parameter inputs) which drew are included. In summary, rotations never appear to perform worse (when evaluated on the secondary outcomes) in the draws as they maintain the resistance at a lower level.

**Table 3:** Comparing Sequences and Rotations.

| Outcome |  | Operational Outcome |  |
| --- | --- | --- | --- |
| Sequence Win | 31119 (14%) | Sequence Operational Win | 15590 (7%) |
| Rotation Win | 30544 (14%) | Rotation Operational Win | 14110 (6%) |
| Draw | 158337 (72%) | No Operational Win | 190300 (87%) |

### Comparing Sequences and Rotations versus Mixtures

Where it is chemically possible to combine two insecticides together into a single mixture insecticide formulation, should these insecticides be deployed as a single mixture formulation or as two separate single insecticide formulations? Mixtures were found never to perform worse when compared against sequences (Figure 6) or rotations (Figure 7) regardless of the starting resistance, deployment interval or degree of cross resistance between the two insecticides when both insecticides were given identical properties.

**Figure 6.**
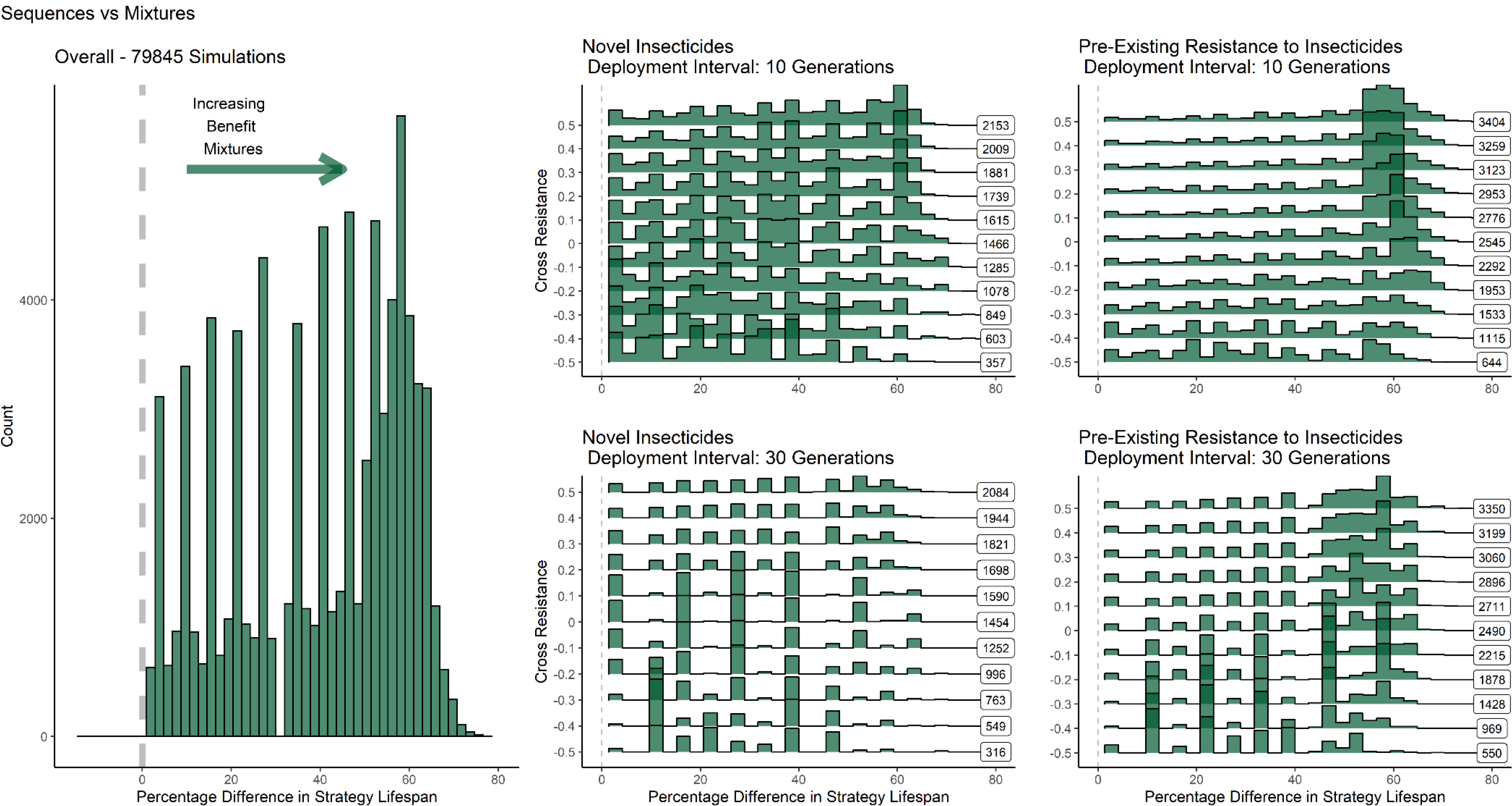
Frequency Distributions Comparing the Strategy Lifespans Between Sequences and Mixtures for Insecticides with Identical Properties. Percentage differences above zero indicate mixtures were the favoured strategy, values below zero favour sequences. Draws (identical strategy lifespans, difference = 0) were excluded from the plots. The left plot is the overall global distribution without any stratification. The four plots on the right are stratified by the deployment interval (top-bottom) and starting resistance (novel: z= 0, pre-used: z = 50) (left-right). Each row in these plots is the frequency distribution of the percentage difference in strategy lifespan for whether cross resistance was positive, negative, or not included. Difference values above 0 (green) favour mixtures. For the cross-selection plots, the numbers on the right-hand side of each plot indicate how many comparisons were in each distribution (that is, the number of comparisons where the simulations did not draw). In summary, mixtures never appear to be inferior to sequences when both insecticides have equivalent properties.

**Figure 7.**
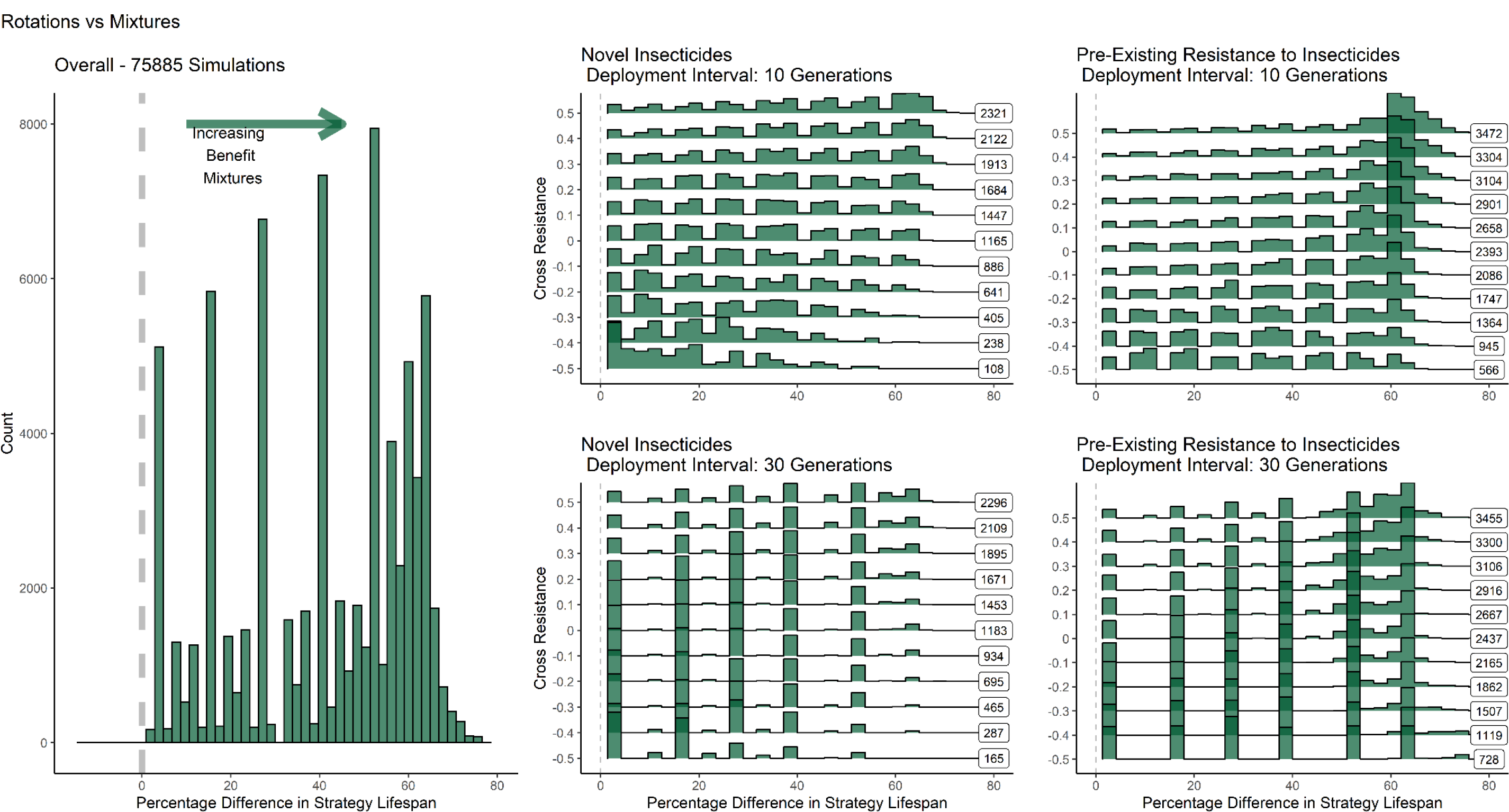
Frequency Distributions Comparing the Strategy Lifespans between Rotations and Mixtures for Insecticides with Identical Properties. This figure has exatly the same structure and interperetion as Figure 6, it simply compares rotations (rather than sequences) agianst mixtures

### Comparing Sequences, Rotations, Adaptive Rotations and Mixtures with Unique Insecticides

When allowing the insecticides to have unique properties (starting resistance, heritability, and fitness cost), the rotation strategy performed badly. However, this is due to the deployment rules of the rotation model, which prevent immediate redeployment i.e., if one insecticide has failed there is no longer anything available to be rotated so the simulation is terminated, meaning that the strategy can no longer be implemented. When the deployment rules are relaxed allowing rotations to be adaptive rotations (defaulting to be in sequence when necessary), adaptive rotations generally outperformed sequences (Figure 8). This is expected because the adaptive rotation strategy rotates but with facility to become a temporary sequence when required.

**Figure 8.**
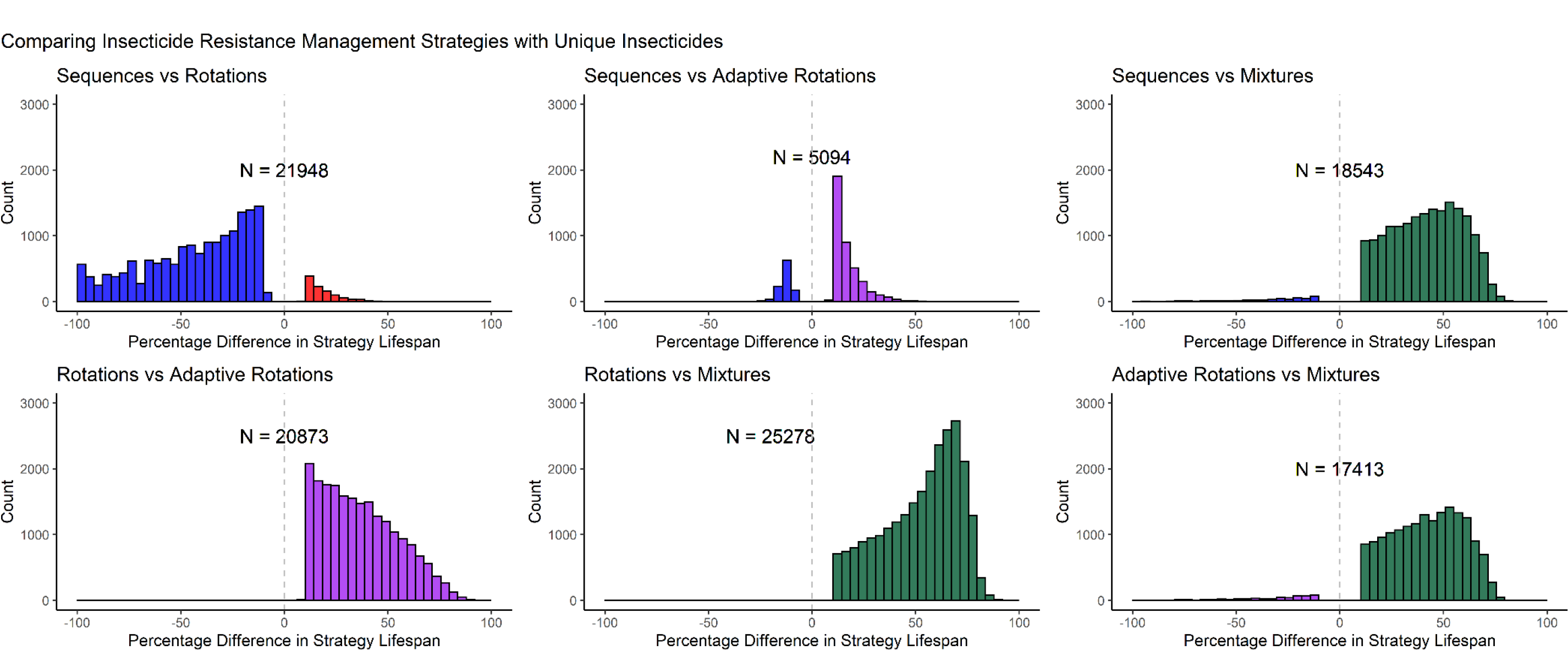
Frequency Distribution of the Percentage Difference in Strategy Lifespan when Insecticides have unique properties. Top-Left: Comparing sequences versus rotations. Top-Middle: Comparing sequences versus adaptive rotations. Top-Right: Comparing sequences versus mixtures. Bottom-left: comparing rotations versus adaptive rotations. Bottom-middle comparing rotations versus mixtures. Bottom-right:comparing adaptive rotations versus mixtures. Plots only include comparisons where the difference in strategy lifespan was ≥10%, where N is the number of comparisons (out of a potential 50,000) included in each plot. Colours represent which strategy won: Blue = Sequences, Red = Rotations, Purple = Adaptive Rotations, Green = Mixutres. Note, the sequence versus rotation plot highlights the issue with the restrictive deployment rules of the rotation strategy when comparing insectcides with unique properties.

When the insecticides in the mixtures were given unique properties (starting resistance, heritability, and fitness cost), there are occasions when mixtures are no longer the dominant strategy, although these situations were rare when compared to the number of mixture wins (Figure 8). In these situations, the insecticides were very dissimilar from one another in terms of their starting PRS and heritability (Supplement 3, Figure S2). This indicates that only one insecticide in the mixture failed, and the other insecticide would remain available to be deployed singularly as a sequence.

### Identifying Parameters Driving Insecticide Resistance to Inform Decision Making

There are situations where while on average two or more IRM strategies may perform equally well (Table 3 and Table 4), but under a narrowed parameter space one strategy is preferential to another. It is important here to identify what conditions lead to such events and which parameters or subset of parameters lead to such outcomes occurring. These are identified using three statistical methods in the subsections below. This can inform what field data would be most valuable to collect to help choose between IRM strategies. Identifying which parameters offer both a good predictive value, while also ideally being simplistic and reliable to measure in the field, would help to improve the choice of IRM strategy.

**Table 4:** Comparing Sequences, Rotations and Mixtures.

| Table 4: Comparing Sequences, Rotations and Mixtures |  |  |  |  |
| --- | --- | --- | --- | --- |
| Outcome |  |  | Operational Outcome |  |
| Draw | 138105<br>(63%) |  | No Operational Win | 146285<br>(66%) |
| Mixture Win | 73835<br>(34%) |  | Mixture Operational Win | 65978<br>(30%) |
| Rotation Loss<br>(Sequence-Mixture Draw) | 2050 (1%) |  | Rotation Loss (Sequence and<br>Mixture Operational Draw) | 2620 (1%) |
| Sequence Loss<br>(Rotation-Mixture Draw) | 6010 (3%) |  | Sequence Loss (Rotation and<br>Mixture Operational Draw) | 5117 (2%) |

### Partial Rank Correlation

Partial rank correlation (Figure 9) looks for correlations between the parameter input value and the outcome measure, in this case the strategy lifespan, and acts as a way of determining which parameters drive or inhibit the evolution of resistance. Increasing values of parameters with negative correlation drive the evolution of IR, while increasing the values of those with positive correlation slow the evolution of IR. The following parameters were identified as driving faster evolution of IR: male insecticide exposure, female insecticide exposure, intervention coverage and heritability. Fitness costs were associated with the slowing of the evolution of IR. Mosquito dispersal had no clear impact on the evolution of IR. These drivers of IR (as quantified by degree of correlation) were independent of cross resistance between insecticides.

**Figure 9.**
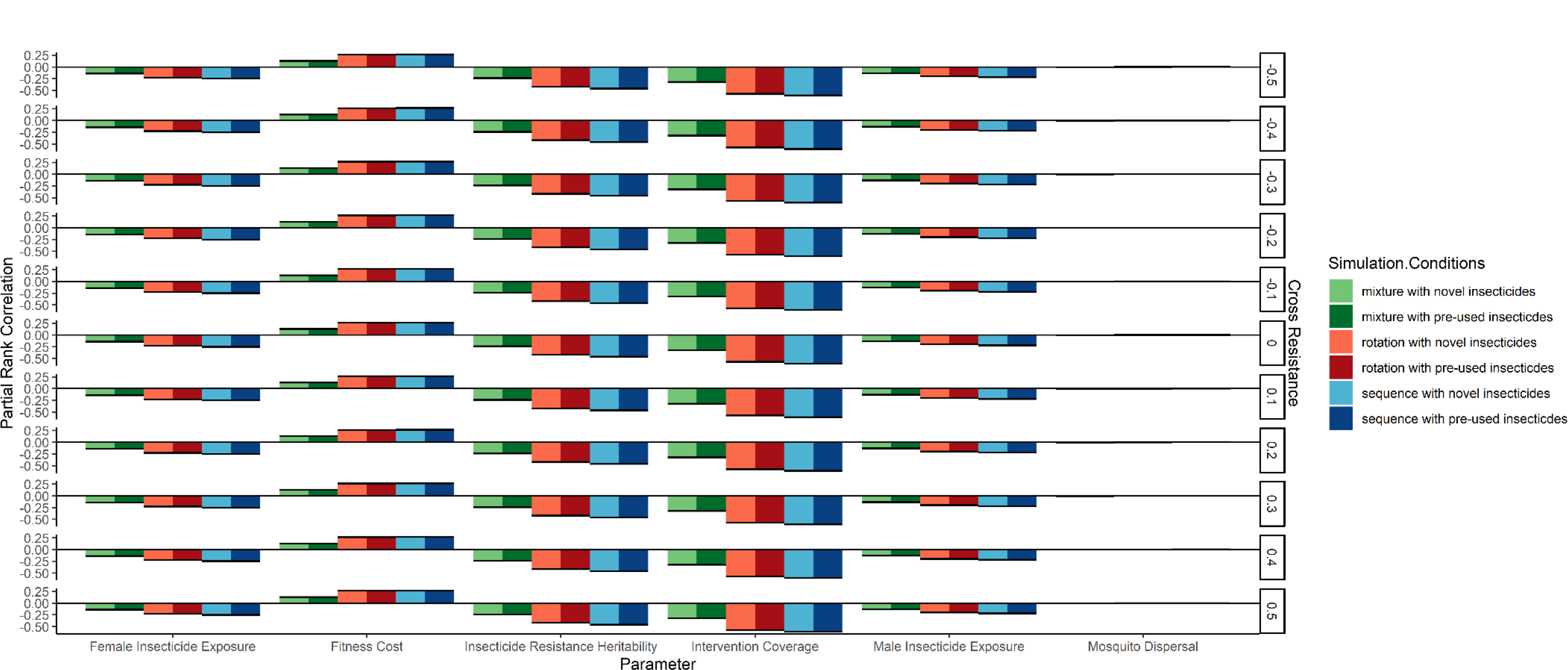
Partial Rank Correlation Between Parameter Inputs and Strategy Lifespan. Partial rank correlation was stratified by the IRM strategy (sequence [blues], rotation [reds], mixture [greens]), the starting polygenic resistance score of the simulation (novel = 0, pre-existing resistance = 50; darker colours indicate the simulations had pre-existing resistance. Each panel is the degree of cross resistance between the two insecticides. Error bars are the 95% confidence intervals of the correlation estimate. Negative estimates are detrimental to IRM (decrease the longevity of the simulation), positive estimates are beneficial to IRM (increase the strategy lifespan).

### Generalised Linear Modelling

Negative estimates indicate the strategy lifespan decreases as the parameter value increases. Positive estimates indicate an increase in the strategy lifespan as the parameter value increases. Mosquito dispersal was found to have a complex non- linear relationship (Supplement 3, Figure S3). Very low and very high rates of dispersal were found to slow the evolution of IR and prolong the strategy lifespans; however, all other rates of dispersal were found to increase the evolution of IR (i.e., to reduce the strategy lifespans). This non-linear relationship helps to explain the near zero correlation obtained for dispersal using partial rank correlation (Figure 4). Heritability was found to have the largest effect size on the strategy lifespan. The effect size differences between sequences and rotations were small when compared against the mixture baseline (Table 5).

**Table 5:**
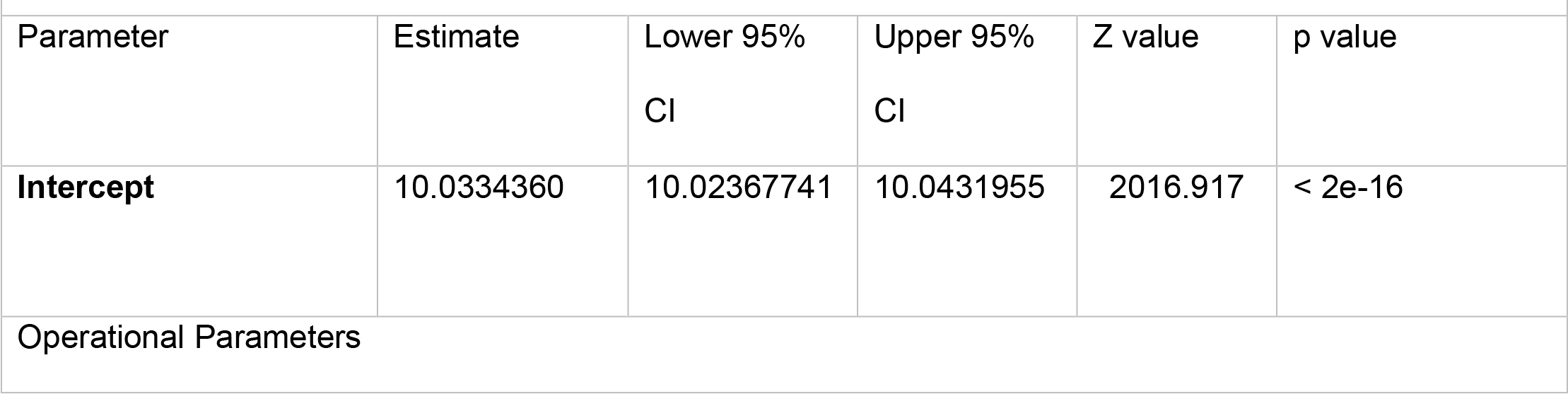
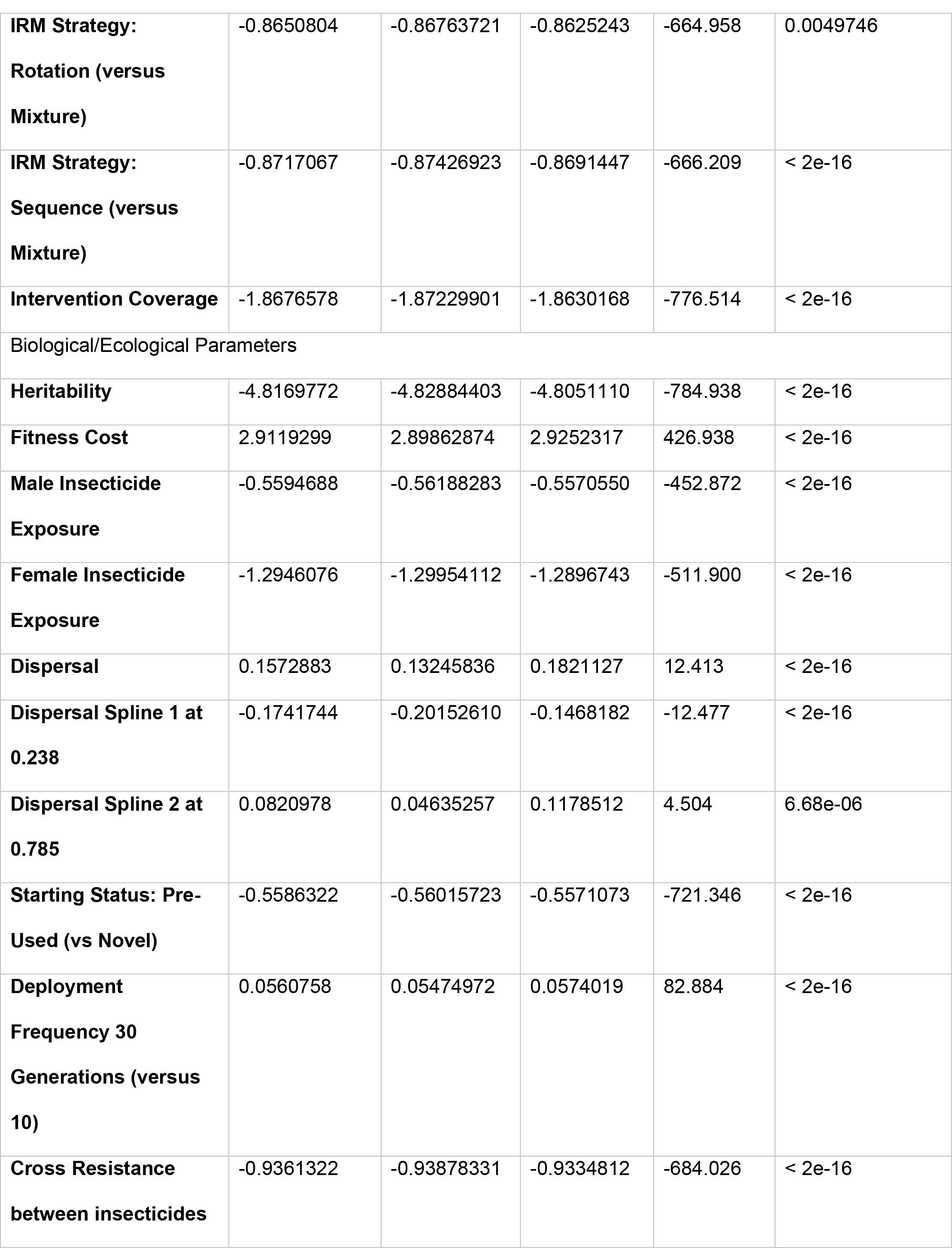
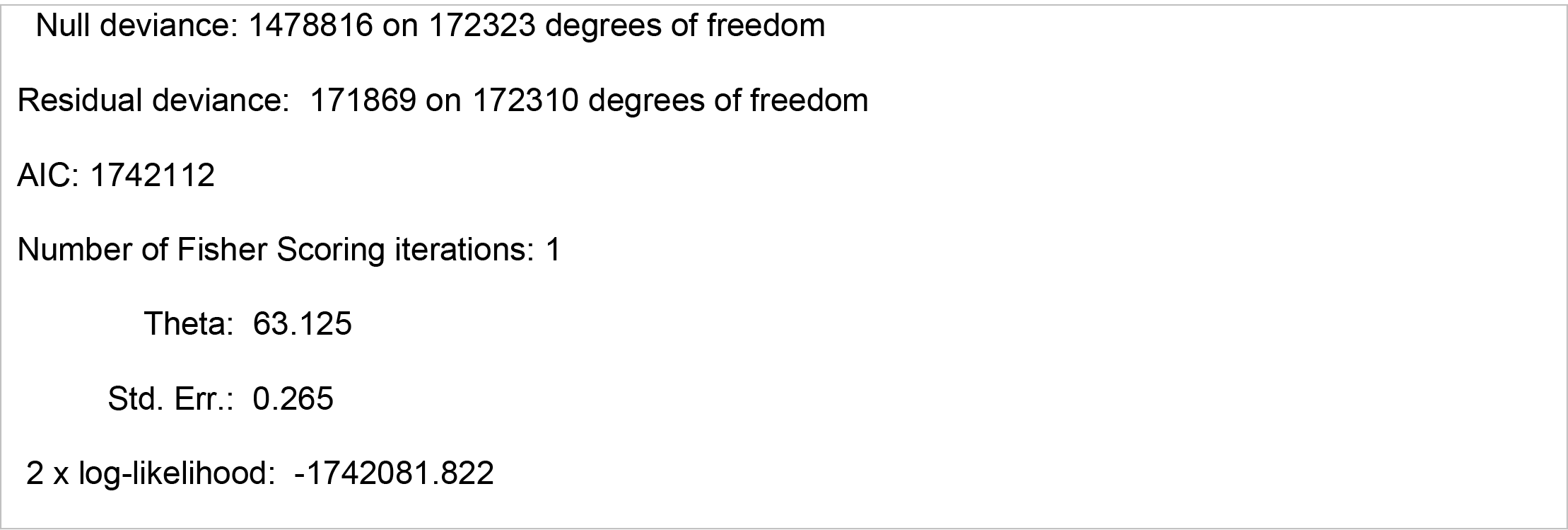
Negative Binomial Generalised Linear Model Output.

### Random Forest Models

A total of eight random forest models were fitted on the operational outcome predicted by the model inputs, with the aim to identify parameters useful for predicting which strategy to use. Random forest models allowed for the identification of parameters which offer predictive power in determining which IRM strategy to use. Parameter importance is identified by the mean decrease in accuracy. Where higher values indicate these parameters are more important in the model, and therefore more important in terms of providing information on which to make informed IRM decisions.

Figure 10 shows the parameter importance when choosing between sequences and rotations. Figure 10A being the parameter importance when both insecticides had equal properties, and highlights intervention coverage and the degree of cross resistance between the insecticides as important predictors. Figure 10B shows the parameter importance when both insecticides were allowed to have unique properties, here intervention coverage and cross resistance were also found to be the predictors with the greatest importance. Figure 11 shows the parameter importance when choosing between sequences, rotations, and mixtures. Figure 11A is for when both insecticides had the same properties and highlights the importance of intervention coverage followed by heritability. Figure 11B is when the insecticides had unique properties intervention coverage became more important (relative to the other predictors), with cross resistance the next most important predictor.

**Figure 10.**
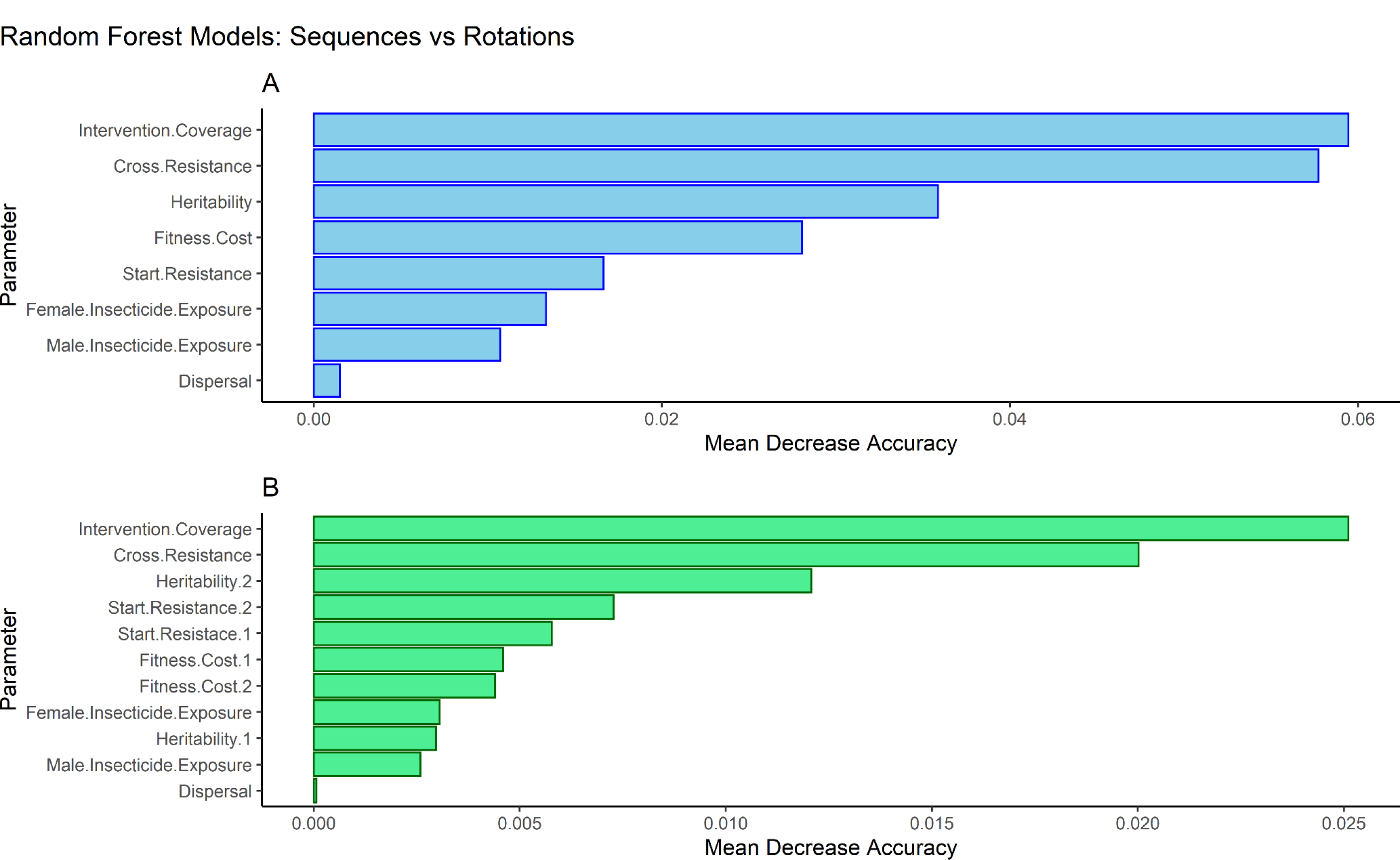
Parameter Importance from Random Forest Model for Choosing between Sequences and Rotations. Panel A (blue bars) is the parameter importance for when both insecticides have equivalent properties and has a predictive accuracy of 87.65%. Panel B (green bars) is the parameter importance when the insecticides are given unique properties and has a predictive accuracy of 90.98%. The x axis shows the decrease in predictive accuracy when the parameter is removed from the analyse so larger values indicate greater predictive importance. Parameters are ordered by importance.

**Figure 11.**
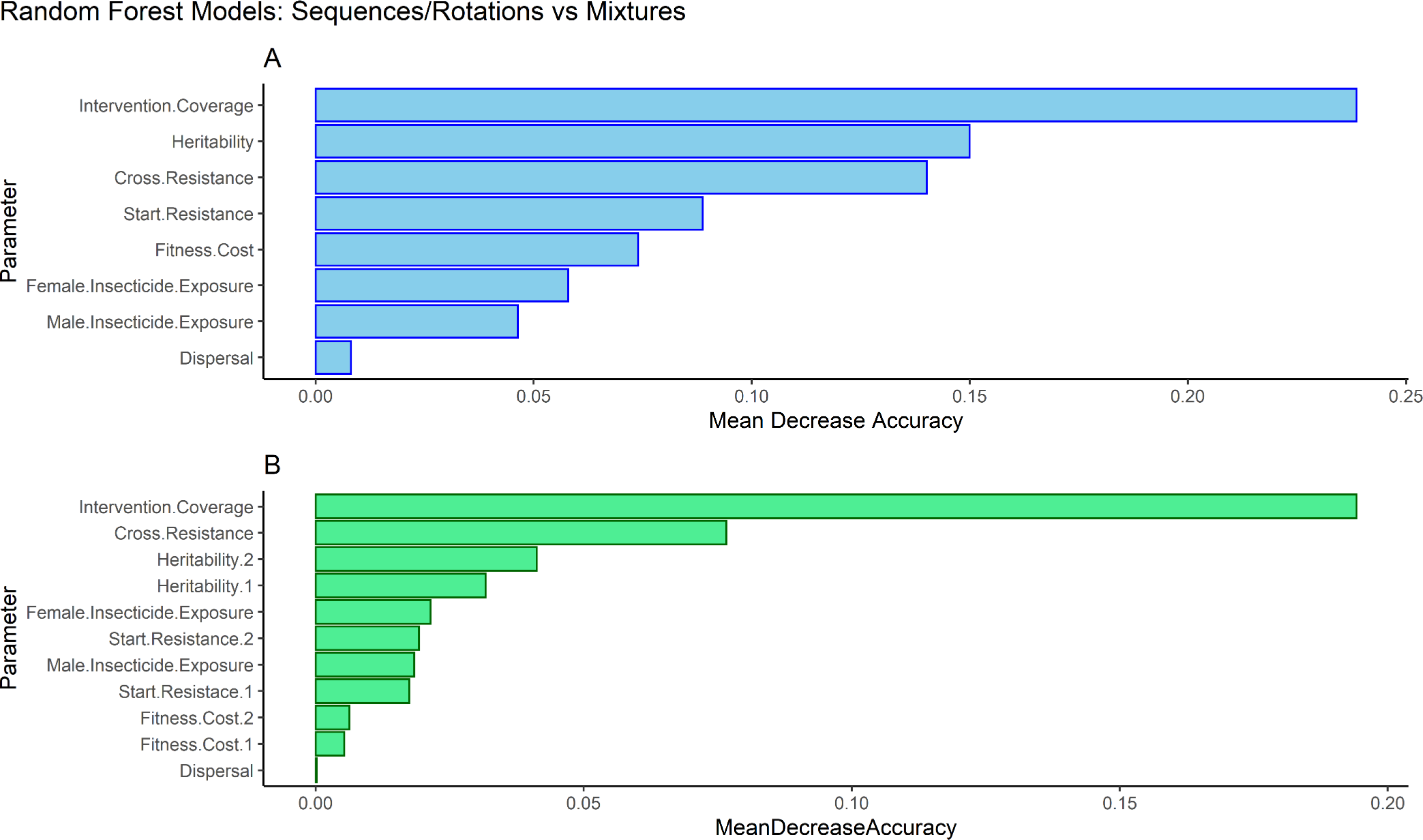
Parameter Importance from Random Forest Model for Choosing between Sequences, Rotations and Mixtures. Panel A (blue bars) is the parameter importance for when both insecticides have identical properties and has a predictive accuracy of 96.14% %. Panel B (green bars) is the parameter importance when the insecticides are given unique properties and has a predictive accuracy of 91.16%. The x axis shows the decrease in predictive accuracy when the parameter is removed from the analyse so larger values indicate greater predictive importance.

Intervention coverage being an important predictor is intuitive because it dictates the amount of insecticide selection. However, its use as a predictor lies more in deciding whether an IRM strategy is needed at all, because at lower intervention coverages, the “no operational win category” is dominant because all strategies are likely to run to the maximum of 500 generations (Figure 12). Therefore, a second set of random forest models was fit, but including only simulations where the intervention coverage was greater than or equal to 0.5, to better identify parameters which help to choose between IRM strategies.

**Figure 12.**
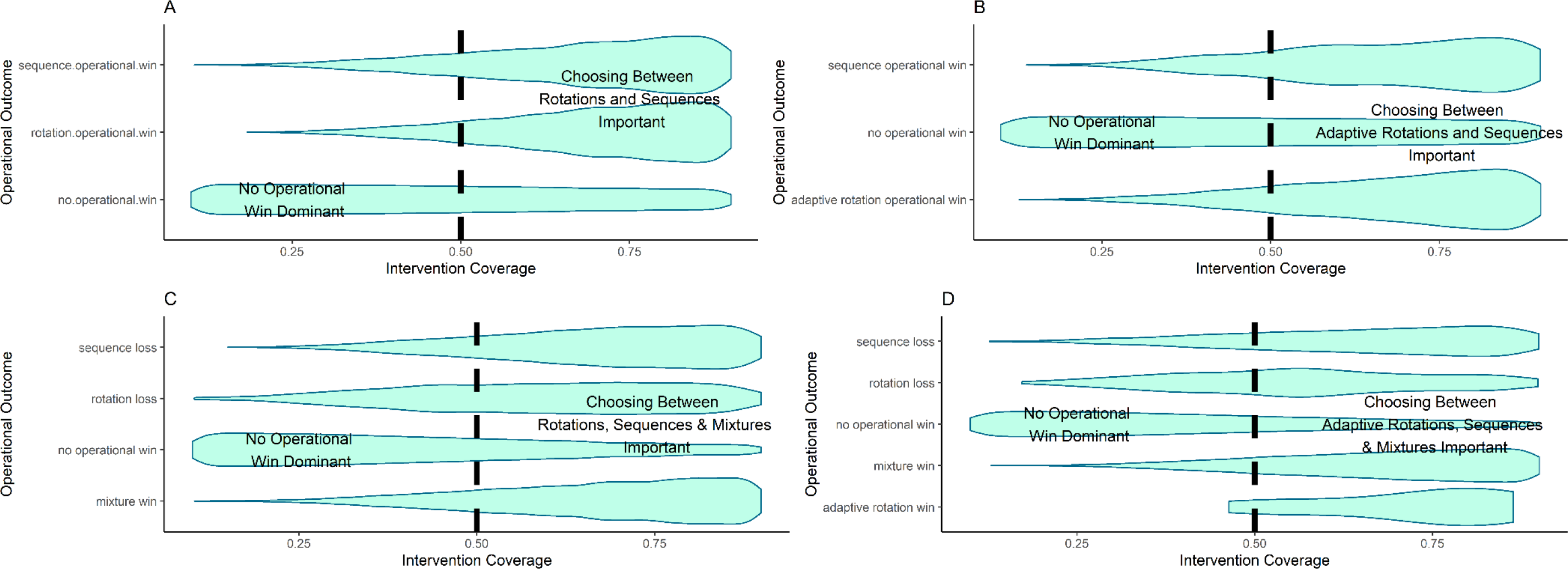
Violin Plot of the Operational Outcome in Relation to the Intervention Coverage. Panel A: Sequences versus Rotations – Equivalent Insecticides. Panel B: Sequences vs Adaptive Rotations – Unique Insecticides. Panel C: Sequences vs Rotations vs Mixtures – Equivalent Insecticides. Pane l D: Sequences vs Adaptative Rotations vs Mixtures – Unique Insecticides. Dotted line is 50% intervention coverage and indicates an approximate point where IRM strategy choice matters, which is the “no operational win” outcome is not dominant.

Figure 13 shows the parameter importance for this random forest model when choosing between sequences and rotations. The intervention coverage parameter is no longer the most important predictor, being replaced by the cross resistance between the insecticides, with heritability and fitness costs also being important predictors for simulations where the insecticides had equal properties (Figures 13A). When allowing the insecticides to have unique properties cross resistance is the most important parameter alongside heritability (Figures 13B). Figure 14 shows the parameter importance for choosing between sequences, rotations, and mixtures. Heritability and cross resistance being important predictors when the insecticides have equal properties (Figures 14A) or unique properties (Figure 14B).

**Figure 13.**
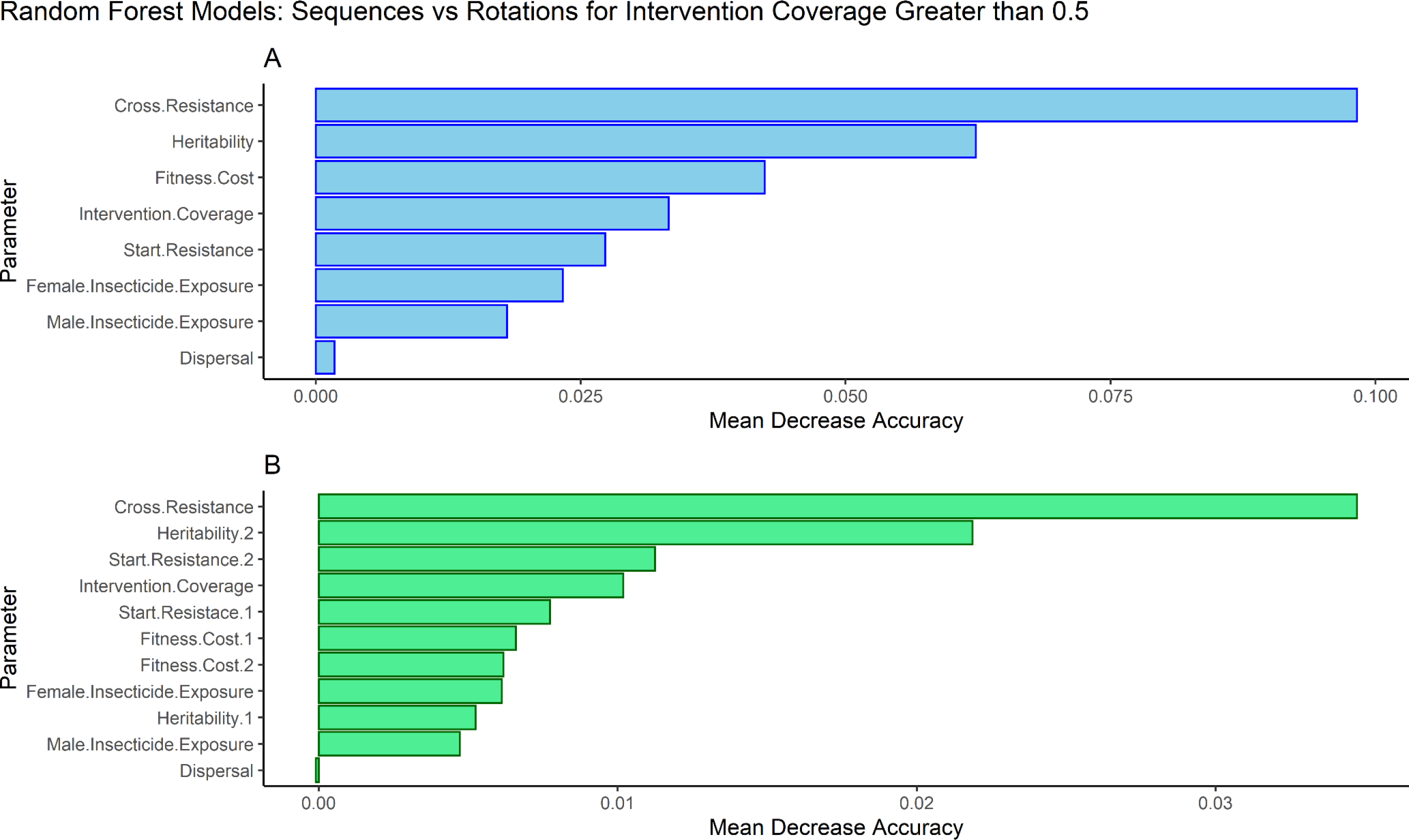
Parameter Importance from Random Forest Model for Choosing between Sequences and Rotations restricted to simulations where Intervention Coverage ≥0.5. Random forest model was restricted to comparisons where intervention coverage ≥0.5. Panel A (blue bars) is the parameter importance for when both insecticides have equivalent properties and has a predictive accuracy of 79.09%. Panel B (green bars) is the parameter importance when the insecticides are given unique properties and has a predictive accuracy of 84.73%. The x axis shows the decrease in predictive accuracy when the parameter is removed from the analyse so larger values indicate greater predictive importance.

**Figure 14.**
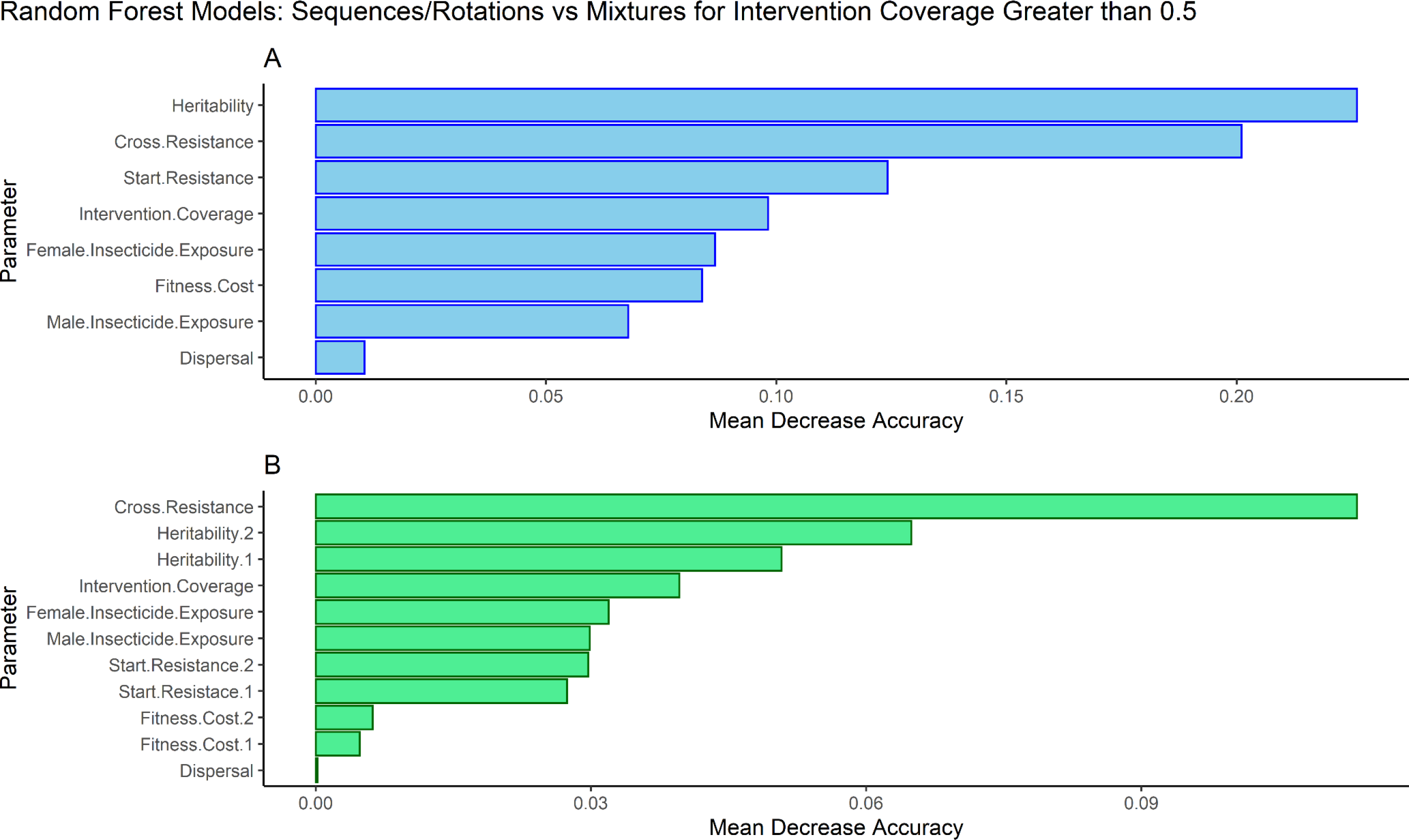
Parameter Importance from Random Forest Models for choosing between sequences, rotations and mixtures restricted to simulations where Intervention Coverage ≥0.5. Panel A (blue bars): Random Forest model parameter importance for simulations where insecticides were given equal parameters, model accuracy 94.32%. Panel B (green bars) : Random Forest model parameter importance for simulations where insecticides were given unique properties, model accuracy 87.63%. The x axis shows the decrease in predictive accuracy when the parameter is removed from the analyse so larger values indicate greater predictive importance.

## Discussion

We modelled polygenic IR as a quantitative trait by devising a Polygenic Resistance Score that can be measured as WHO cylinder bioassay survival. Previous models have applied a quantitative genetics framework to IR traits (Gardner et al., 1998; Haridas & Tenhumberg, 2018), but did not investigate the implications for resistance management strategies utilising multiple insecticides. Gardner et al looked at the impact of different dosing strategies on the rate of the evolution of polygenic resistance, a tactic not readily available in public health where uniform insecticide dosing is intended through the use or LLINs or IRS. Utilising quantitative genetics has allowed for a conceptually simplistic method for incorporating cross resistance/selection into our model, a process generally absent from monogenic models (Rex Consortium, 2010). We used genetic correlations allowing us to explore the possible implications of cross resistance for IRM. A headline summary of the results from all the tables and figures can be found in Table 6. There are of course important caveats of the model to address to provide context to our results.

**Table 6:** Summary of Results Tables and Figures.

| Table 6: Summary of Results Tables and Figures |  |
| --- | --- |
|  | Take Home Message(s) / Purpose |
| Tables |  |
| 1 | Summary of all model assumptions |
| 2 | Summary of all parameter inputs |
| 3 | Rotations perform marginally better than sequences. |
| 4 | Mixtures perform much better than rotations and sequences. |
| 5 | Sequence and Rotation effect size similar, smaller than mixtures. |
| Figures |  |
| 1 | Polygenic Resistance Score (PRS) measured as bioassay survival. |
|  | Bioassay survival converted to expected field survival. |
| 2 | The model is calibrated such that an “average” insecticide lasts 10 years. |
| 3 | Explains how insecticides are deployed and the concept of the withdrawal and return thresholds. |
| 4 | Sequences and Rotations perform equally at the “global level”<br>Rotation better than sequences negative cross resistance.<br>Sequences slightly better than rotations with positive cross resistance. |
| 5 | When sequences and rotations draw on strategy lifespan rotations have:<br>• Lower peak bioassay survival<br>• Lower mean bioassay survival to deployed insecticide. |
| 6 | Mixtures outperform sequences regardless of cross resistance. |
| 7 | Mixtures outperform rotations regardless of cross resistance. |
| 8 | When insecticides have unique properties:<br>• Mixtures perform best<br>• Rotations perform worst<br>• Adaptive rotations best solo deployment strategy. |
| 9 | Drivers of Resistance (in order of importance):<br>• Intervention Coverage<br>• Heritability<br>• Female Insecticide Exposure<br>• Male Insecticide Exposure |
| 10 | Highest importance variables for choosing between sequences and rotations:<br>• Intervention Coverage<br>• Cross Resistance<br>• Heritability<br>• Fitness Cost |
| 11 | Highest importance variables for choosing between sequences, rotations, and mixtures:<br>• Intervention coverage<br>• Heritability |
|  | • Cross Resistance<br>• Start Resistance |
| 12 | Intervention coverage predominantly determines if IRM is necessary. |
| 13 | Highest importance variables for choosing between sequences and rotations:<br>• Cross resistance<br>• Heritability<br>• Fitness Cost<br>• Start Resistance |
| 14 | Highest importance variables for choosing between sequences, rotations, and mixtures:<br>• Heritability<br>• Cross Resistance<br>• Intervention Coverage |

### Model Caveats

The first caveats concern the rules of insecticide deployment and withdrawal used. We highlight our simulations were terminated under idealistic deployment conditions when mosquito survival is still relatively low (i.e., <10% in bioassays). Setting the withdrawal threshold higher (e.g., 20, 30 or 50% bioassay survival), would inevitably lead to more strategies drawing (as they would run to 500 generation completion), and requiring comparison based only on the secondary outcomes. Or setting a higher withdrawal threshold would require re-calibrating the simulations such that the exposure scaling factor is increased also to maintain an average 10-year insecticide lifespan under continuous deployment.

Insecticide deployments in the models are based on clearly defined pre-set rules required for implementation in computing (see Table 1 for a detailed summary of the model assumptions). In practice, deployments are often affected by logistical factors such as unreliable supply chains, human resource limitations, and slow, centralised decision making. The primary outcome of strategy lifespan does not readily allow for suitable comparison between different deployment intervals. This is because the deployment interval dictates the frequency at which the model can be interrogated and stopped if an IRM strategy has failed (i.e., if both insecticides are above the withdrawal threshold of 10% bioassay survival). If both insecticides reach the 10% bioassay threshold in ten-generation deployment interval five generations after deployment, then the simulation terminates five generations later. However, if the deployment interval was 30 generations, the simulation would continue for a further 25 generations before terminating, artificially inflating the apparent strategy lifespan. We feel this reflects the operational reality as LLINs are designed for 3-year deployments (∼30 generations) and IRS for 1-year deployments (∼10 generations) and it is unlikely that detection of resistance would result in replacement of nets after 1.5 years or IRS after 6 months. Insecticides are also deployed “perfectly” in our models, and we do not account for economic constraints dictating the continued use of cheaper insecticides. This is especially important when considering mixtures, as these formulations are expected to be more expensive and there will be an economic temptation to reduce the concentration of each insecticide in the mixture. Neither do we allow for any logistical constraints with regards to insecticide deployment. Insecticide deployments always occur instantaneously at the set time (without the delays often noted in practice). Insecticide choices are also made “perfectly”, the resistance in the field is tracked without error and information is available to make instantaneous, correct decisions regarding insecticide deployment. There is likely to be a considerable lag between insecticide susceptible testing, insecticide procurement and insecticide deployment resulting from inadequate surveillance systems and the release of procurement funds (Chanda et al., 2015). In operational reality, insecticides are highly unlikely to be deployed in the idealistic ways the rules the model demands. It is also unlikely that a single IRM strategy would remain fixed for eternity, with an amalgamation of sequences, rotations and mixtures potentially being used in a single intervention site. We could only realistic investigate the IRM strategies under these idealised conditions of “perfect” deployment because space restraints prevented a more extended investigation of how operation factors may alter the relative merits (although the model can later be used to investigate specific examples of operational problems). We can however speculate on how operation limitations may affect the strategies. Mixtures are likely to be the most robust because no replacement decisions are required (Figure 3) supporting our conclusion that full-dose mixtures are overall the “best” IRM strategy. Sequences and rotations performed very similarly in our simulations but as noted by Curtis (1987) and Hastings et al., (2022). the fact that rotations have pre-planned insecticide replacement already in place may make them more operationally robust than sequences whose unpredictable replacement intervals require rapid responses.,

Second are caveats concerning non-public health insecticide selection pressure. The model only tracks insecticides in the armoury and assumes these insecticides are only ever deployed (and therefore encountered by mosquitoes) as part of a public heath deployment. Other insecticides, which mosquitoes may encounter such as agricultural insecticides or personal household insecticides (i.e., coils and aerosols) were not considered in the model as they are not part of the armoury and we assumed they had no cross resistance with insecticides in the armoury. Agricultural insecticide use is associated with increased IR in malaria vectors (Abuelmaali et al., 2013; Nkya et al., 2014; Reid & McKenzie, 2016). This further extends to the model’s intervention site and refugia existing “isolated” from rest of the world. It would be possible to build these “external” selection pressures into our model (through Equations 3a and 3b) and to allow them to act even in refugia (by adding an additional term representing external selection in Equation 9a) but this is a specific operational question we could not realistically address in this study due to time constraints.

### Model Interpretation

With these caveats in mind, we can say, from a purely IRM perspective the overall best strategy appears to be full-dose mixtures (Figures 6 and 7). This broadly corroborates with the results from monogenic models (e.g. Madgwick & Kanitz, 2022; South & Hastings, 2018) for favouring full-dose mixtures. There appears to be little difference between rotations and sequences when comparing operational longevity at the “global level” (Figure 4) agreeing with monogenic modelling results (Hastings et al., 2022), although rotations are slightly more beneficial when considering mean and peak bioassay survival (Figure 5). The reason for this would appear to be that sequences are in practice a type of rotation Hastings et al (2022) i.e., because the first insecticide in the sequence can later be re-deployed if resistance has fallen below its redeployment threshold). A blend of these two strategies (which we defined as adaptive rotations) generally was the better strategy when considering solo insecticide deployments (Figure 8).

These conclusions ignore important operational and economic factors required for the IRM implementation, for example the increased costs of a mixture LLIN/IRS, the infrastructure and supply chains required for annual rotations, and so on. However, these results highlight a vital conclusion from our study i.e., the underlying assumption of a purely monogenic or purely polygenic basis of IR does not overly impact the general conclusions from mathematical models; lending further support to mixtures being the generally optimal IRM strategy. Validating IRM models with field data is challenging, probably impossible, because of the need for long field surveillance of the levels of resistance the need to measure other parameters associated with quantifying the degree of selection, and the need for replication over several IRM sites to draw robust empirical conclusions. Therefore, mathematical modelling studies using different approaches and assumptions yet yielding qualitatively similarly conclusions suggests there is some robustness to the conclusions made.

One advantage of taking a quantitative genetics approach is the ability to easily incorporate cross resistance/selection using correlated responses. One important concern with mixtures has been the role cross resistance could play (e.g Curtis, 1985). Importantly, the presence of cross resistance between insecticides in the mixture does not appear to invalidate the conclusion that mixtures are the more beneficial IRM strategy (Figures 6 and 7), despite previous speculation cross-resistance would compromise mixtures as an IRM strategy. There are two possible explanations for this. First, because cross resistance also reduces the strategy lifespan of sequences and rotations through indirect selection occurring on the non-deployed insecticide. Second, our mixture consists of both insecticides at full-dose and this double-dose effect likely overcomes the detrimental impacts of cross resistance. In the few examples where a monogenic model has included cross resistance this pattern was also observed (Birch & Shaw, 1997; Roush, 1998; Sudo et al., 2018). This is certainly not to say we (nor indeed the authors of the previous studies) recommend mixing two insecticides where cross-resistance or cross-selection is likely, but that if these two insecticides were to be deployed in the same area mixtures would very likely be a better IRM strategy deploying them singularly as sequences or rotations. We would instead emphasise the importance of insecticides in the entire insecticide armoury being chosen that have as a little cross-resistance as possible. Note, the simulations were terminated when mosquito survival was still relatively low (i.e., <10% in bioassays). At higher resistance levels mosquito survival increases, and the protective effect of each mixture component therefore decreases. Whether cross-resistance becomes more detrimental under lower insecticide doses and higher resistances needs investigation, as these are both real-world problems.

Mathematical modelling allows insights into how decisions could or should be made from a policy or operational perspective, and also identifies important research and knowledge gaps through the identification of understudied yet operationally important variables. Our analysis highlights the importance of cross resistance between insecticides, intervention coverage, IR heritability and the fitness costs associated with IR as being important predictors for which IRM strategy to deploy (Figure 14). Unfortunately, the heritability of resistance is challenging to reliably measure in the laboratory (Rosenheim, 1991) and, in any case, such estimates are unlikely to reflect heritability in the field where environmental variance will be much higher. Fitness costs are also challenging, as these can affect a wide range of life-history traits and are difficult to measure outside unrealistic laboratory settings. There is also large variation in the design and execution of fitness studies (Freeman et al., 2021) due to the large number of traits measured. Our model included the fitness costs as a single global value to account for cumulative detrimental effects fitness costs can play across the whole life-history of a mosquito.

For malaria, the primary purpose for deploying insecticides is reducing its transmission and the associated burden of disease rather than managing IR in the mosquito population. The choice of IRM strategy should of course not be conducted at the detriment of inadequate malaria control. For example, removing the insecticide selection pressure entirely would allow for any fitness costs to return the population to susceptibility but in doing so there is no mosquito control and therefore no disease control. But at the same time, immediate reductions in malaria transmission (to meet a particular policy objective) without considering the longer-term sustainability of such insecticidal deployment could lead to the loss of future transmission reductions and the resurgence of malaria transmission. There is a need to manage the longer-term effectiveness of insecticides alongside the immediate epidemiological benefits of deploying insecticides and we argue that computer simulation is the most feasible way to identify best-practice in deployment of insecticides used for vector control.

## Acknowledgements

We thank Andy South for comments on early drafts and coding advice

## Funding

NH is an MRC-DTP candidate. The funder had no role in the study design, data collection, interpretation of data, writing of paper or decision to publish.

## Competing Interests

The authors declare they have no competing interests.

## Data Availability and Code

Model code for running and analysing simulations is available from https://github.com/NeilHobbs/polyres and/or the corresponding author on request.

## Supplement 1: Linear modelling between bioassay survival and field survival

### Bioassay Survival and Field Survival Conversion

Equation 1b requires the conversion of the tracked bioassay survival (based on the Polygenic Resistance Score) to a field survival proportion. The relationship between bioassay survival and field survival was estimated using matched WHO cylinder bioassays with experimental hut trial data (data obtained from Churcher et al. 2016). In this study Churcher et al. used compared the link between bioassay mortality and experimental hut mortality for LLINs. For the quantitative genetics model the relationship between bioassay survival and experimental hut survival is needed, rather than the relationship between mortalities. The obtained mortality data (both experimental hut and bioassay) was therefore converted to survival data. Generalised additive modelling was used to visually confirm the linear relationship. We used data obtained from by Churcher et al (2016) but restricted our model to only include the insecticide deltamethrin and *An. gambiae* s.l. mosquitoes, to reduce any confounding effects between different species responses to different insecticides. The total number of observations used was therefore 13. A simple linear model was then fit to the proportion surviving in the experimental hut against the proportion dying in the WHO cylinder:

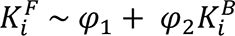

This can be used to estimate the relationship between bioassay survival and expected field survival. For the model, we use the estimate of 0.15 for the intercept and 0.48 for the regression coefficient, such that both values are rounded to two decimal places (Table S1).

**Table S1:** Linear Model of Field Survival and Bioassay Survival.

| Table S1: Linear Model of Field Survival and Bioassay Survival |  |  |  |  |  |
| --- | --- | --- | --- | --- | --- |
| Parameter | Estimate | Upper 95% CI | Lower 95% CI | t value | P value |
| Intercept | 0.14581 | 0.3649152 | -0.07329093 | 1.465 | 0.1710 |
| Bioassay Survival | 0.48438 | 0.7949830 | 0.17376870 | 3.432 | 0.0056 |
| Residual standard error: 0.1644 on 11 degrees of freedom |  |  |  |  |  |
| Multiple R-squared: 0.5171, Adjusted R-squared: 0.4732 |  |  |  |  |  |
| F-statistic: 11.78 on 1 and 11 DF, p-value: 0.0056 |  |  |  |  |  |

**Figure S1.**
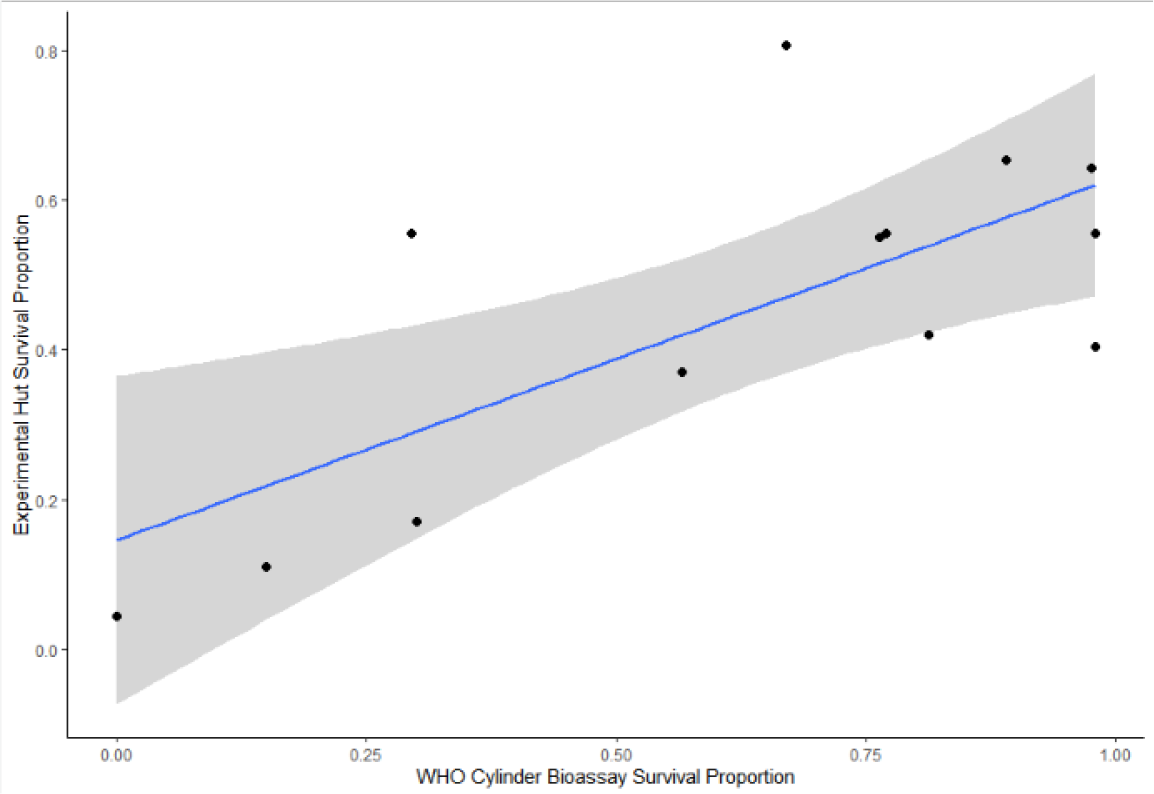
Relationship between experimental hut survival and bioassay survival. Regression line with 95% CI.

### Supplement S2: Stability of the Polygenic Resistance Score with Population Standard Deviation

**Table S2:**
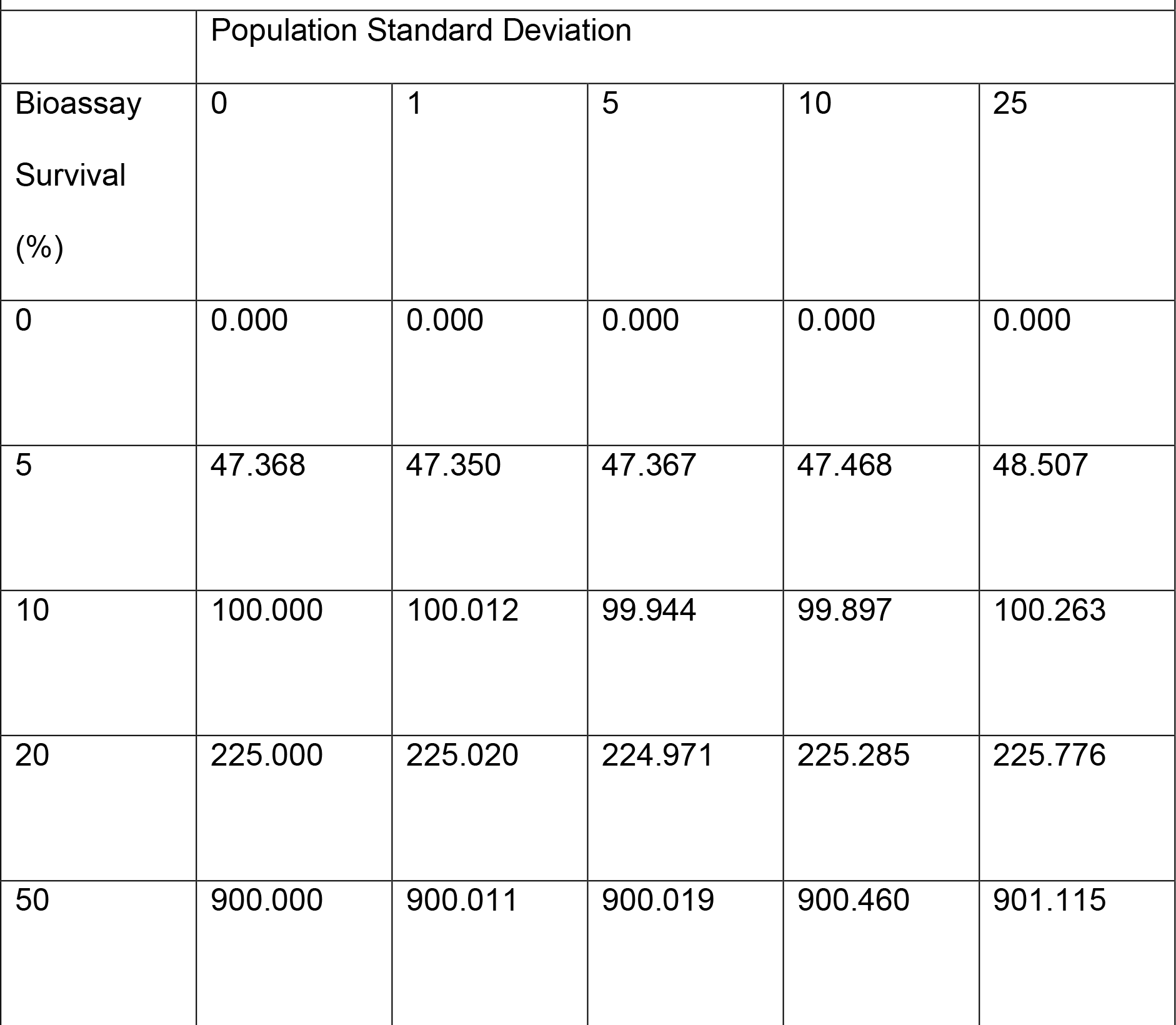
Stability of the Polygenic Resistance Score with Population Standard Deviation.

### Supplement 3: Additional Exploratory and Descriptive Plots

**Figure S2.**
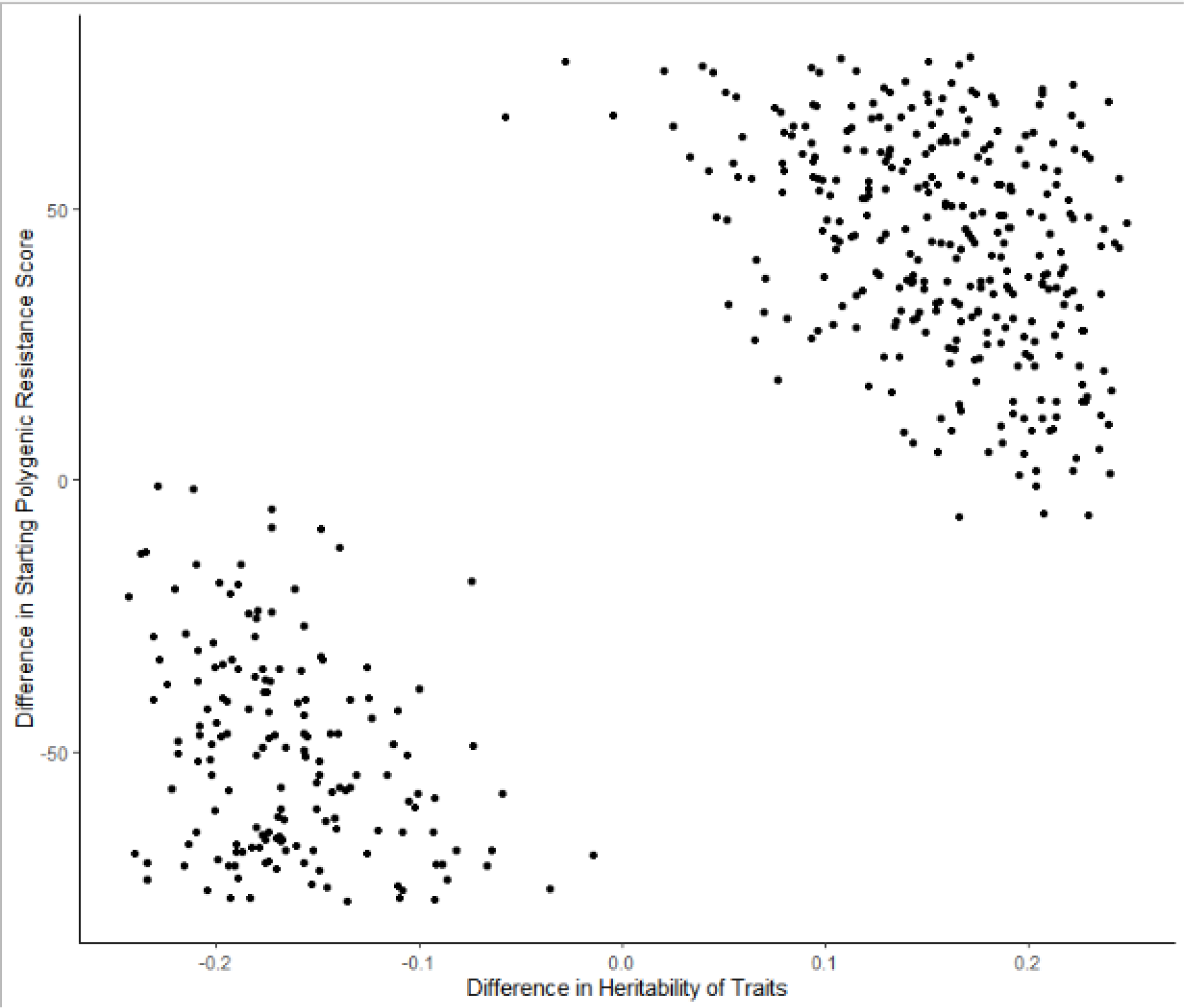
Explaining Scenarios where Sequences outperformed Mixtures. Highlights that for mixtures to be effective as an IRM strategy need to have broadly equal rates of evolution (equal heritability of traits) and broadly equal starting resistance values.

**Figure S3.**
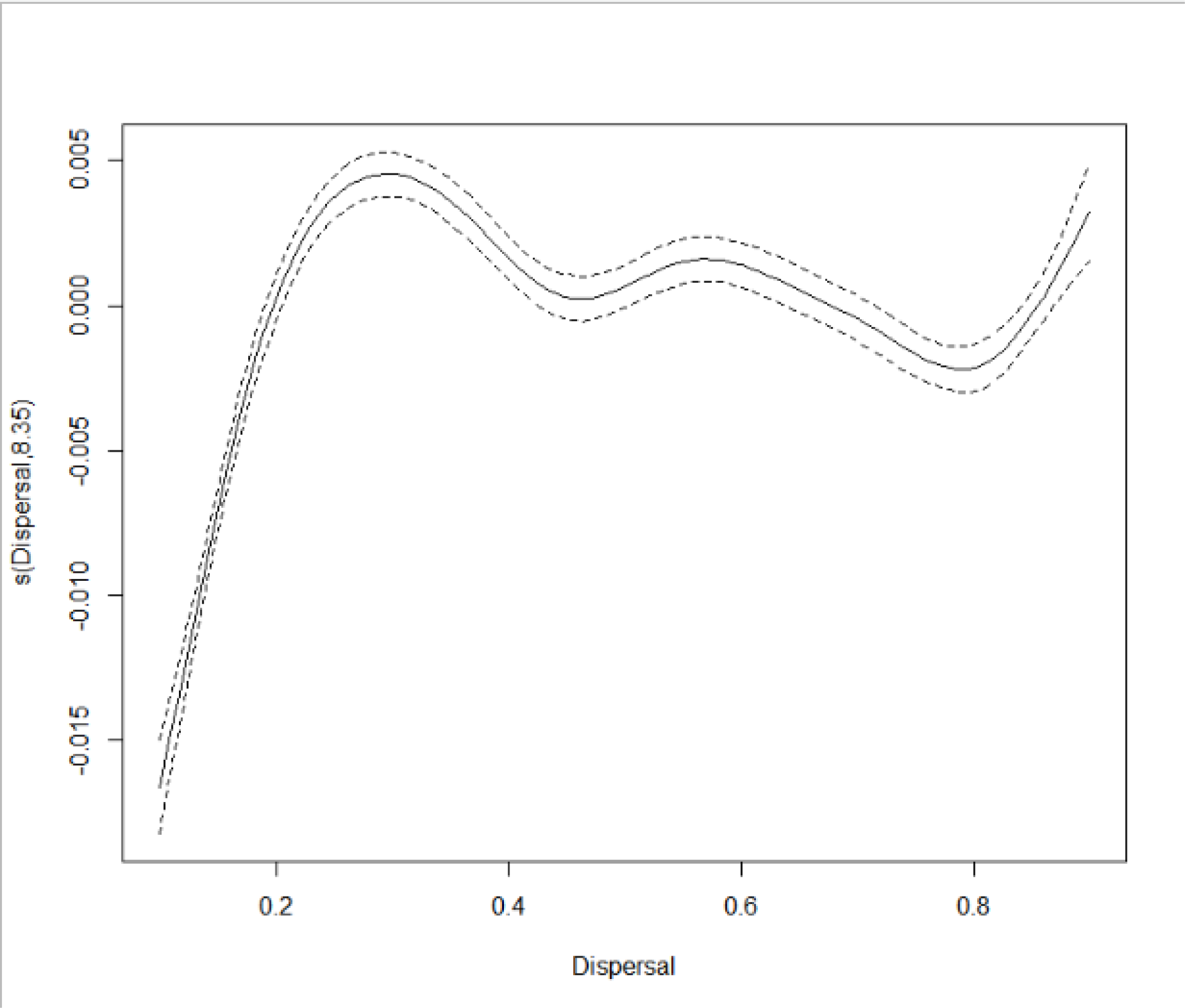
Generalised Additive Model smoothing over Dispersal to identify knot positions for piecewise GLM. Where the x-axis the dispersal rate, and the y-axis is the smoothing of the Dispersal parameter, and the effective degrees of freedom. High y-axis values indicate the simulation duration would be increased, while lower y-axis values indicate the simulation duration would be decreased. This plot was simply used to identify the knot positions of the dispersal parameter for the GLM.

## Notes

### Competing Interest Statement

The authors have declared no competing interest.

## References

1. Abuelmaali, S. A., Elaagip, A. H., Basheer, M. A., Frah, E. A., Ahmed, F. T. A., Elhaj, H. F. A., Seidahmed, O. M. E., Weetman, D., & Hamid, M. M. A. (2013). Impacts of agricultural practices on insecticide resistance in the malaria vector Anopheles arabiensis in Khartoum state, Sudan. PLoS ONE, 8(11), 1–9. https://doi.org/10.1371/journal.pone.0080549

2. Asidi, A., N’Guessan, R., Akogbeto, M., Curtis, C., & Rowland, M. (2012). Loss of household protection from use of insecticide-treated nets against pyrethroid- resistant mosquitoes, Benin. Emerging Infectious Diseases, 18(7), 1101–1106. https://doi.org/10.3201/eid1807.120218

3. Balabanidou, V., Grigoraki, L., & Vontas, J. (2018). Insect cuticle: a critical determinant of insecticide resistance. Current Opinion in Insect Science, 27(Figure 1), 68–74. https://doi.org/10.1016/j.cois.2018.03.001

4. Bhatt, S., Weiss, D. J., Cameron, E., Bisanzio, D., Mappin, B., Dalrymple, U., Battle, K. E., Moyes, C. L., Henry, A., Eckhoff, P. A., Wenger, E. A., Briët, O., Penny, M. A., Smith, T. A., Bennett, A., Yukich, J., Eisele, T. P., Griffin, J. T., Fergus, C. A., … Gething, P. W. (2015). The effect of malaria control on Plasmodium falciparum in Africa between 2000 and 2015. Nature, 526(7572), 207–211. https://doi.org/10.1038/nature15535

5. Birch, C. P. D., & Shaw, M. W. (1997). When can Reduced Doses and Pesticide Mixtures Delay the Build-up of Pesticide Resistance? A Mathematical Model. The Journal of Applied Ecology, 34(4), 1032. https://doi.org/10.2307/2405292

6. Brooke, B. D. (2008). Kdr: Can a Single Mutation Produce an Entire Insecticide Resistance Phenotype? Transactions of the Royal Society of Tropical Medicine and Hygiene, 102(6), 524–525. https://doi.org/10.1016/j.trstmh.2008.01.001

7. Carnell, R. (2020). *lhs: Latin Hypercube Samples* (R package version 1.0.2). https://cran.r-project.org/package=lhs

8. Chanda, E., Mzilahowa, T., Chipwanya, J., Mulenga, S., Ali, D., Troell, P., Dodoli, W., Govere, J. M., & Gimnig, J. (2015). Preventing malaria transmission by indoor residual spraying in Malawi: Grappling with the challenge of uncertain sustainability. Malaria Journal, 14(1), 1–7. https://doi.org/10.1186/s12936-015-0759-3

9. Churcher, T. S., Lissenden, N., Griffin, J. T., Worrall, E., & Ranson, H. (2016). The impact of pyrethroid resistance on the efficacy and effectiveness of bednets for malaria control in Africa. ELife, 5(AUGUST), 1–26. https://doi.org/10.7554/eLife.16090

10. Comins, H. N. (1977). The development of insecticide resistance in the presence of migration. Journal of Theoretical Biology, 64(1), 177–197. https://doi.org/10.1016/0022-5193(77)90119-9

11. Curtis, C. F. (1985). Theoretical models of the use of insecticide mixtures for the management of resistance. Bulletin of Entomological Research, 75(2), 259–266. https://doi.org/10.1017/S0007485300014346

12. Curtis, C. (1987). Genetic aspects of selection for resistance. In M. G. Ford, D. W. Hollomon, & B. P. S. Khambay (Eds.), Combating resistance to xenobiotics: Biological and chemical approaches. Ellis Horwood.

13. Freeman, J. C., Smith, L. B., Silva, J. J., Fan, Y., Sun, H., & Scott, J. G. (2021). Fitness studies of insecticide resistant strains: lessons learned and future directions. *Pest Management Science*, January. https://doi.org/10.1002/ps.6306

14. Gardner, S. N., Gressel, J., & Mangel, M. (1998). A revolving dose strategy to delay the evolution of both quantitative vs major monogene resistances to pesticides and drugs. International Journal of Pest Management, 44(3), 161–180. https://doi.org/10.1080/096708798228275

15. Grossman, M. K., Oliver, S. V., Brooke, B. D., & Thomas, M. B. (2020). Use of alternative bioassays to explore the impact of pyrethroid resistance on LLIN efficacy. Parasites and Vectors, 13(1), 1–12. https://doi.org/10.1186/s13071-020-04055-9

16. Hancock, P. A., Hendriks, C. J. M., Tangena, J. A., Gibson, H., Hemingway, J., Coleman, M., Gething, P. W., Cameron, E., Bhatt, S., & Moyes, C. L. (2020). Mapping trends in insecticide resistance phenotypes in African malaria vectors. PLoS Biology, 18(6), 1–23. https://doi.org/10.1371/journal.pbio.3000633

17. Haridas, C. V., & Tenhumberg, B. (2018). Modeling effects of ecological factors on evolution of polygenic pesticide resistance. Journal of Theoretical Biology, 456, 224–232. https://doi.org/10.1016/j.jtbi.2018.07.034

18. Hastings, I., Jones, S., & South, A. (2022). Evaluating rotations as a strategy to delay the evolution of insecticide resistance in vectors of human diseases . 1–38. https://doi.org/10.1101/2022.10.07.511276

19. Helps, J. C., Paveley, N. D., & van den Bosch, F. (2017). Identifying circumstances under which high insecticide dose increases or decreases resistance selection. Journal of Theoretical Biology, 428, 153–167. https://doi.org/10.1016/j.jtbi.2017.06.007

20. Hemingway, J., Ranson, H., Magill, A., Kolaczinski, J., Fornadel, C., Gimnig, J., Coetzee, M., Simard, F., Roch, D. K., Hinzoumbe, C. K., Pickett, J., Schellenberg, D., Gething, P., Hoppé, M., & Hamon, N. (2016). Averting a malaria disaster: Will insecticide resistance derail malaria control? The Lancet, 387(10029), 1785–1788. https://doi.org/10.1016/S0140-6736(15)00417-1

21. Irish, S., N’Guessan, R., Boko, P., Metonnou, C., Odjo, A., Akogbeto, M., & Rowland, M. (2008). Loss of protection with insecticide-treated nets against pyrethroid-resistant Culex quinquefasciatus mosquitoes once nets become holed: An experimental hut study. Parasites and Vectors, 1(1), 1–5. https://doi.org/10.1186/1756-3305-1-17

22. Kwiatkowska, R. M., Platt, N., Poupardin, R., Irving, H., Dabire, R. K., Mitchell, S., Jones, C. M., Diabaté, A., Ranson, H., & Wondji, C. S. (2013). Dissecting the mechanisms responsible for the multiple insecticide resistance phenotype in Anopheles gambiae s.s., M form, from Vallée du Kou, Burkina Faso. Gene, 519(1), 98–106. https://doi.org/10.1016/j.gene.2013.01.036

23. Levick, B., South, A., & Hastings, I. M. (2017). A Two-Locus Model of the Evolution of Insecticide Resistance to Inform and Optimise Public Health Insecticide Deployment Strategies. In PLoS Computational Biology (Vol. 13, Issue 1). https://doi.org/10.1371/journal.pcbi.1005327

24. Liaw, A., & Wiener, M. (2002). Classification and Regression by randomForest. R News, 2, 18–22.

25. Lindsay, S. W., Thomas, M. B., & Kleinschmidt, I. (2021). Threats to the effectiveness of insecticide-treated bednets for malaria control: thinking beyond insecticide resistance. The Lancet Global Health, 9(9), e1325–e1331. https://doi.org/10.1016/S2214-109X(21)00216-3

26. Lissenden, N., Kont, M. D., Essandoh, J., Ismail, H. M., Churcher, T. S., Lambert, B., Lenhart, A., McCall, P. J., Moyes, C. L., Paine, M. J. I., Praulins, G., Weetman, D., & Lees, R. S. (2021). Review and meta-analysis of the evidence for choosing between specific pyrethroids for programmatic purposes. Insects, 12(9), 1–22. https://doi.org/10.3390/insects12090826

27. Madgwick, P. G., & Kanitz, R. (2022a). Beyond redundant kill : a fundamental explanation of how insecticide mixtures work for resistance management Running title : Beyond redundant kill This article is protected by copyright . All rights reserved . Abstract. Pest Management Science. https://doi.org/10.1002/ps.7180

28. Madgwick, P. G., & Kanitz, R. (2022b). Modelling new insecticide-treated bed nets for malaria-vector control: how to strategically manage resistance? Malaria Journal, 21(1), 1–20. https://doi.org/10.1186/s12936-022-04083-z

29. Mani, G. S. (1985). Evolution of resistance in the presence of two insecticides. Genetics, 109(4), 761–783. https://doi.org/10.1093/genetics/109.4.761

30. Nkya, T. E., Poupardin, R., Laporte, F., Akhouayri, I., Mosha, F., Magesa, S., Kisinza, W., & David, J. P. (2014). Impact of agriculture on the selection of insecticide resistance in the malaria vector Anopheles gambiae: A multigenerational study in controlled conditions. Parasites and Vectors, 7(1), 1–12. https://doi.org/10.1186/s13071-014-0480-z

31. Oliver, S. V., & Brooke, B. D. (2013). The effect of larval nutritional deprivation on the life history and DDT resistance phenotype in laboratory strains of the malaria vector Anopheles arabiensis. Malaria Journal, 12(1), 1–9. https://doi.org/10.1186/1475-2875-12-44

32. Osoro, J. K., Machani, M. G., Ochomo, E., Wanjala, C., Omukunda, E., Munga, S., Githeko, A. K., Yan, G., & Afrane, Y. A. (2021). Insecticide resistance exerts significant fitness costs in immature stages of Anopheles gambiae in western Kenya. Malaria Journal, 20(1), 1–7. https://doi.org/10.1186/s12936-021-03798-9

33. Oxborough, R. M. (2016). Trends in US President’s Malaria Initiative-funded indoor residual spray coverage and insecticide choice in sub-Saharan Africa (2008- 2015): Urgent need for affordable, long-lasting insecticides. Malaria Journal, 15(1), 1–9. https://doi.org/10.1186/s12936-016-1201-1

34. Peck, S. L. (2001). Antibiotic and insecticide resistance modeling - Is it time to start talking? Trends in Microbiology, 9(6), 286–292. https://doi.org/10.1016/S0966-842X(01)02042-X

35. R Core Team. (2020). R: A language and environment for statistical computing. (4.0.3). R Foundation for Statistical Computing. https://www.r-project.org/

36. Ranson, H., & Lissenden, N. (2016). Insecticide Resistance in African Anopheles Mosquitoes: A Worsening Situation that Needs Urgent Action to Maintain Malaria Control. Trends in Parasitology, 32(3), 187–196. https://doi.org/10.1016/j.pt.2015.11.010

37. Ranson, H., N’Guessan, R., Lines, J., Moiroux, N., Nkuni, Z., & Corbel, V. (2011). Pyrethroid resistance in African anopheline mosquitoes: What are the implications for malaria control? Trends in Parasitology, 27(2), 91–98. https://doi.org/10.1016/j.pt.2010.08.004

38. Reid, M. C., & McKenzie, F. E. (2016). The contribution of agricultural insecticide use to increasing insecticide resistance in African malaria vectors. Malaria Journal, 15(1), 1–8. https://doi.org/10.1186/s12936-016-1162-4

39. Rex Consortium. (2007). Structure of the scientific community modelling the evolution of resistance. PLoS ONE, 2(12). https://doi.org/10.1371/journal.pone.0001275

40. Rex Consortium. (2010). The skill and style to model the evolution of resistance to pesticides and drugs. Evolutionary Applications, 3(4), 375–390. https://doi.org/10.1111/j.1752-4571.2010.00124.x

41. Rex Consortium. (2013). Heterogeneity of selection and the evolution of resistance. Trends in Ecology and Evolution, 28(2), 110–118. https://doi.org/10.1016/j.tree.2012.09.001

42. Rosenheim, J. A. (1991). Realized heritability estimation for pesticide resistance traits. Entomologia Experimentalis et Applicata, 58(1), 93–97. https://doi.org/10.1111/j.1570-7458.1991.tb01456.x

43. Roush, R. T. (1998). Two-toxin strategies for management of insecticidal transgenic crops: Can pyramiding succeed where pesticide mixtures have not? Philosophical Transactions of the Royal Society B: Biological Sciences, 353(1376), 1777–1786. https://doi.org/10.1098/rstb.1998.0330

44. South, A., & Hastings, I. M. (2018). Insecticide resistance evolution with mixtures and sequences: A model-based explanation. Malaria Journal, 17(1), 1–20. https://doi.org/10.1186/s12936-018-2203-y

45. Stevenson, M., Nunes, T., Heuer, C., Marshall, J., Sanchez, J., Thorn-Ton, R., Reiczigel, J., Robison-Cox, J., Sebastiani, P., Solymos, P., Yoshida, K., Jones, G.-O., Pirikahu, S., & Maintainer, S. F. (2020). *Tools for the Analysis of Epidemiological Data* (R package version 2.0.17.). https://cran.r-project.org/package=epiR

46. Sudo, M., Takahashi, D., Andow, D. A., Suzuki, Y., & Yamanaka, T. (2018). Optimal management strategy of insecticide resistance under various insect life histories: Heterogeneous timing of selection and interpatch dispersal. Evolutionary Applications, 11(2), 271–283. https://doi.org/10.1111/eva.12550

47. Tabashnik, B. E. (1986). Computer simulation as a tool for pesticide resistance management. Pesticide Resistance: Strategies and Tactics for Management, 195–203.

48. Tchouakui, M., Riveron Miranda, J., Mugenzi, L. M. J. J., Djonabaye, D., Wondji, M. J., Tchoupo, M., Tchapga, W., Njiokou, F., & Wondji, C. S. (2020). Cytochrome P450 metabolic resistance (CYP6P9a) to pyrethroids imposes a fitness cost in the major African malaria vector Anopheles funestus. Heredity, 124(5), 621– 632. https://doi.org/10.1038/s41437-020-0304-1

49. Van Hul, N., Braks, M., & Van Bortel, W. (2021). A systematic review to understand the value of entomological endpoints for assessing the efficacy of vector control interventions. June, 36.

50. Venables, W. N., & Ripley, B. D. (2002). Modern Applied Statistics with S (Fourth Edi). Springer.

51. Via, S. (1986). Quantitative genetic models and the evolution of pesticide resistance. Pesticide Resistance: Strategies and Tactics for Management, 222–235.

52. Walsh, B., & Lynch, M. (2018). Evolution and Selection of Quantitative Traits. Oxford University Press.

53. WHO. (2012). Global Plan for Insecticide Resistance Managment.

54. WHO. (2018). Test procedures for insecticide resistance monitoring in malaria vector mosquitoes (Second edition) (Updated June 2018). In *Who*. http://www.who.int/malaria/publications/atoz/9789241511575/en/

55. WHO. (2020). Preferred Product Characteristics : insecticide-treated nets for malaria transmission control in areas with insecticide-resistant mosquito populations Draft for public consultation.

56. Wickham, H. (2009). ggplot2: Elegant Graphics for Data Analysis. Springer-Verlag. http://ggplot2.org

57. Wilson, A. L., Boelaert, M., Kleinschmidt, I., Pinder, M., Scott, T. W., Tusting, L. S., & Lindsay, S. W. (2015). Evidence-based vector control? Improving the quality of vector control trials. Trends in Parasitology, 31(8), 380–390. https://doi.org/10.1016/j.pt.2015.04.015

58. Wood, R. J. (1981). Strategies for conserving susceptibility to insecticides. Parasitology, 82, 69–80.

59. Wood, S. N. (2011). Fast stable restricted maximum likelihood and marginal likelihood estimation of semiparametric generalized linear models. Journal of the Royal Statistical Society. Series B: Statistical Methodology, 73(1), 3–36. https://doi.org/10.1111/j.1467-9868.2010.00749.x

